# Loss of MTPAP disrupts mitochondrial RNA processing causing upregulation of type I interferon signalling

**DOI:** 10.64898/2026.05.04.722669

**Authors:** Alexandre Pierga, Yan de Souza Angelo, Adham Safieddine, Marie-Noelle Benassy, Blaise Didry-Barca, Marie Rozen, Luís Seabra, Claire Lovo, Sylvie Bannwarth, Sandra Lacas-Gervais, Gittan Kollberg, Carola Hedberg-Oldfors, Antri Savvidou, Dominique Weil, Yanick J. Crow, Alice Lepelley

## Abstract

Mitochondrial poly-A polymerase (MTPAP) is essential for mitochondrial mRNA (mt-mRNA) polyadenylation, a critical step in mt-mRNA maturation. Mutations in *MTPAP* have been reported to cause a mitochondrial cytopathy. Here, we provide evidence of enhanced type I interferon (IFN) signalling in the blood of patients carrying mutations in *MTPAP*. Further, deletion of *MTPAP* in a fibroblast cell model led to abnormalities in mitochondrial respiration and mtRNA processing, and an upregulation of type I IFN signalling. Notably, both in patients fibroblast and in MTPAP-deleted cells, we observed the accumulation of non-coding mtRNA and mitochondrial double-stranded RNA (mt-dsRNA) within enlarged mitochondrial RNA granules. Cytosolic release of mt-dsRNA led to type I IFN induction mediated primarily by the RNA sensor MDA5 and its adaptor MAVS. Our findings reveal a novel consequence of MTPAP dysfunction, highlighting how impaired mtRNA maturation can drive innate immune system activation.

## INTRODUCTION

Mitochondria are double-membrane, endosymbiotic organelles essential for energy production, metabolic regulation and innate immunity (Suomalainen & Nunnari, 2024). Cells contain multiple copies of the mitochondrial circular genome, with mitochondrial DNA (mtDNA) transcribed bidirectionally from the heavy and light strands into polycistronic transcripts. These transcripts are subsequently cleaved into mature mitochondrial mRNA (mt-mRNA) encoding 13 protein subunits of the oxidative phosphorylation (OXPHOS) complexes, two ribosomal RNAs (mt-rRNA) and 22 transfer RNAs (mt-tRNA) (Chrzanowska-Lightowlers & Lightowlers, 2024), with most of these transcripts derived from the mtDNA heavy strand. Importantly, the majority of proteins involved in mitochondrial processes, including mtDNA replication, transcription, mtRNA maturation, processing and translation, are nuclear-encoded and imported into the organelle (Gustafsson, Falkenberg & Larsson, 2016).

mtRNA maturation takes place within mitochondrial RNA granules (MRGs), which are frequently located adjacent to mtDNA nucleoids (Iborra et al., 2004; Antonicka et al., 2013; Jourdain et al., 2013). These dynamic and fluid compartments, associated with the mitochondrial inner membrane, are enriched in proteins such as GRSF1, and members of the FASTK family required for mtRNA processing, maturation and mitoribosome assembly (Xavier & Martinou, 2021; Rey et al., 2020). mtRNA turnover within mitochondria is mediated predominantly by the degradosome, a complex located close to MRGs composed of the helicase SUV3 (encoded by *SUPV3L1*) and the polynucleotide phosphorylase PNPase (encoded by *PNPT1*) (Borowski et al., 2013; Xavier et al., 2024). The degradosome is essential for the elimination of noncoding mtRNA so as to prevent the accumulation of mitochondrial double-stranded RNA (mt-dsRNA) formed by complementary heavy and light strand transcripts (Mai & Lightowlers, 2017; Pearce et al., 2017; Hensen et al., 2019).

It is increasingly recognised that mitochondrial damage and loss of mitochondrial integrity can lead to mtDNA and mtRNA release into the cytosol, with subsequent triggering of innate immune responses (Lepelley et al., 2021a; Rai & Fessler, 2025). Indeed, type I interferon (IFN) signalling is induced upon detection of nucleic acids in the cytosol by pattern recognition receptors such as cGAS, which detects DNA (Sun et al., 2013), and MDA5 or RIG-I recognizing, respectively, long or short 5’ triphosphate dsRNA, inducing the upregulation of IFNβ and IFN-stimulated genes (ISGs) (Rehwinkel et al., 2020). While IFN signalling is essential for defence against viral infection, inappropriate detection of self-nucleic acid by innate sensors can be pathogenic, as exemplified by the type I interferonopathies, monogenic diseases characterised by chronic activation of IFN signalling (Crow & Casanova, 2024).

We have described enhanced type I IFN signalling in patients with mitochondrial disease due to mutations in *PNPT1* and *ATAD3A*, implicating the cytosolic sensing of released mt-dsRNA (PNPT1) and mtDNA (ATAD3A) (Dhir et al., 2018; Lepelley et al. 2021b). Recently, in addition to PNPase, loss of function of several proteins involved in mtRNA metabolism, such as REXO2 and N6AMT1, has been linked to mt-dsRNA accumulation in mitochondria and upregulation of type I IFN signalling, indicating a broader connection between mtRNA processing and engagement of an innate immune response (Idiiatullina et al., 2024; Foged et al., 2025).

Mitochondrial poly-A polymerase, MTPAP, encoded by *MTPAP*, is a noncanonical poly-A polymerase that catalyses the post-transcriptional polyadenylation of mt-mRNAs, contributing to their stability and translation after cleavage of the polycistronic mtRNA (Tomecki et al., 2004; Nagaike et al., 2005; Chujo et al., 2012). *MT-ND6* is the only non-polyadenylated mt-mRNA. Of note, MTPAP has also been reported to polyadenylate mt-tRNAs, thereby contributing to their degradation by the degradosome (Pearce et al., 2017; Tompuu et al., 2018). To date, the consequences of a loss of mtRNA polyadenylation have remained unclear. Notably, *MTPAP* knock-out (KO) in *Drosophila* results in dsRNA accumulation, similar to the effect of *PNPT1* or *SUPV3L1* KO (Pajak et al., 2019). This observation contradicts the expectation of decreased stability and loss of mt-mRNA based upon a proposed role of the poly-A tail as a stabilising factor (Wilson et al., 2014).

The importance of MTPAP in cellular homeostasis is evidenced by genetic disease, with mutations in *MTPAP* identified in patients presenting a range of clinical features including autosomal recessive spastic ataxia type 4 (SPAX4), optic atrophy, developmental delay, cerebellar atrophy, encephalopathy and progressive neuropathy (Crosby et al., 2010; Al-Shamsi et al., 2016; de Souza et al., 2017; Van Eyck et al., 2020; Hiramatsu et al., 2022; Epstein et al., 2024; Ravanbod et al., 2025). Twelve disease-causing *MTPAP* variants in 17 patients have been reported to date, ten of which are located within the highly conserved enzymatic fingers domain (Ravanbod et al., 2025). Loss of poly-A tails on mt-mRNA due to defective MTPAP activity in patient fibroblasts has been shown to lead to a decrease in mt-mRNA levels, and a loss of mitochondrial complex I and IV subunits, resulting in impaired OXPHOS activity and mitochondrial dysfunction (Wilson et al., 2014).

Prompted by the observation of enhanced type I IFN signalling in two patients with mutations in *MTPAP,* here we show that loss of MTPAP in human fibroblasts leads to an accumulation of mt-dsRNA in mitochondria, and their release into the cytosol triggering type I IFN signalling. By investigating the origin of these mt-dsRNA, we demonstrate a critical role for MTPAP in the removal of non-coding mtRNA. Our data highlight MTPAP as a factor essential to prevent deleterious sensing of mitochondrial nucleic acids.

### Loss of MTPAP results in defective mitochondrial respiration associated with an upregulation of type I IFN signalling

Enhanced expression of ISGs was observed on two occasions in the sister of patient P1, with both siblings manifesting a lethal encephalopathy attributed to a homozygous MTPAP mutation (c.1153A>T; p.Ile385Phe) (Table1) (Van Eyck et al., 2020) (although the sister did not undergo diagnostic genotyping). The IFN scores in blood of the sister were 3.5 (decimal age 0.12 years) and 23.0 (decimal age 0.35 years), both exceeding the normal threshold (< 2.466). Similarly, we detected ISGs upregulation in the blood of a previously unreported patient (P2) presenting with ataxia and progressive microcephaly, who carried a homozygous MTPAP variant (c.245G>A; p.Cys82Tyr) (Table1). The IFN score recorded for this patient was 4.3 (age 4.18 years), also above the normal limit (< 2.758).

While further patient material was unavailable from P1, reduced levels of MTPAP protein were observed by western blot in primary fibroblasts from P2 (Supp Figure 1A), consistent with the data in cells from three previously published patients carrying mutations in *MTPAP,* P3, P4 and P5 (Figure 1A, Table 1). P3 and P4 are two siblings affected with a perinatal encephalopathy, reminiscent of the type I interferonopathy Aicardi-Goutières syndrome (AGS) (Van Eyck et al., 2020). They are compound heterozygous for c.1283T>C (p.Ile428Thr) and c.1567C>T (p.Arg523Trp) mutations in *MTPAP*, associated with impaired mtRNA polyadenylation. Patient P5 carries the homozygous missense mutation c.1432A>G (p.Asn478Asp) leading to strongly decreased polyadenylation activity as measured in vitro (Wilson et al., 2014).

**Figure 1:**
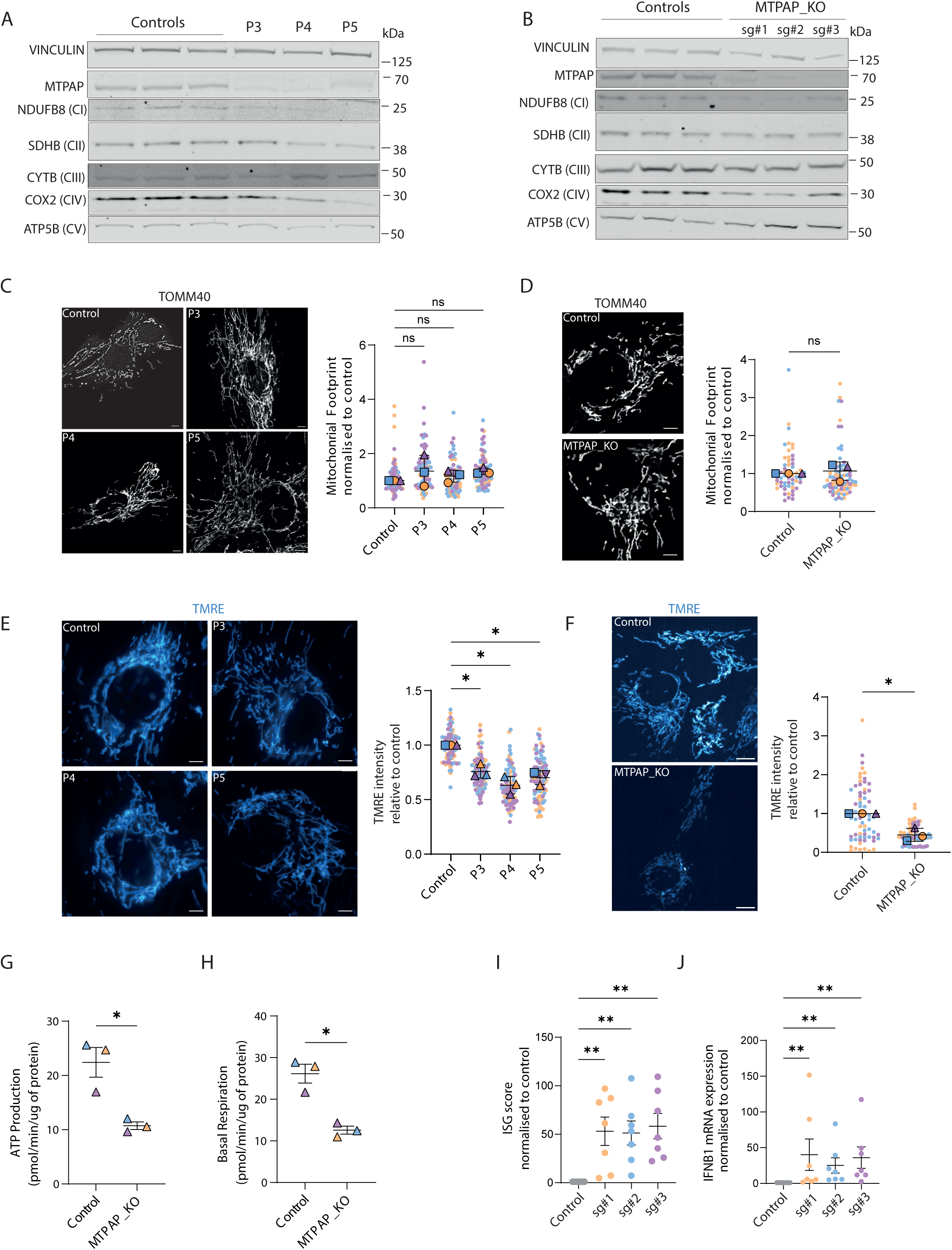
Loss of MTPAP results in defective mitochondrial respiration and upregulation of type I IFN signalling. **A-B:** Western blot showing levels of MTPAP and mitochondrial complex I (CI) NDUFB8, II (CII) SDHB, III (CIII) CYTB, IV (CIV) COX2, V (CV) ATP5B proteins in lysates of fibroblasts from patient P3, P4 and P5 compared to fibroblasts from 3 healthy donor (controls) (**A**), and in lysates of BJ-5ta deleted for *MTPAP* by CRISPR/Cas9 using 3 different single guides (sg#1, #2, #3) compared to controls (**B**). Vinculin is used as a loading control. Image representative of 3 different experiments. **C: Left,** representative confocal images of the mitochondrial network stained with mitochondrial outer membrane protein TOMM40 antibody in patient fibroblasts (P3, P4 and P5) and one healthy donor fibroblast (control). Scale bar: 5 µm. **Right,** quantification of the mitochondrial footprint based on *MiNA* ImageJ analysis plugin in patient fibroblasts and the average of 3 healthy donor fibroblasts (control), represented as a superplot (each border free-point represents the measure of a cell, each colour is a different experiment). Mean ± SEM of 3 experiments; *ns* indicates non significance in a paired t-test performed on the average value for each experiment. **D:** As in C for MTPAP_KO (sg#2) and control cells. **E: Left,** representative spinning disk live images of TMRE signal in patient fibroblasts (P3, P4 and P5) compared to one healthy donor fibroblast (control). Scale bar: 5 µm. **Right,** quantification of TMRE signal in patient fibroblasts relative to the average of 3 healthy donor fibroblasts (control). Superplot, mean ± SEM; n=3 experiments (each colour is a different experiment); each border-free point represents the measure of a cell; * indicates p<0.05 in paired t-test performed on the average value for each experiment. **F:** As in E for MTPAP_KO (sg#2) and control cells. **G-H:** ATP production (**G**) and basal respiration (**H**) measured by Seahorse mito stress assay in MTPAP_KO (average of sg#1, sg#2 and sg#3 data for each experiment) versus control cells. mean ± SEM; n=3 experiments, each colour is a different experiment, * indicates p<0.05 in Mann-Whitney test. **I-J:** ISG score (median of fold changes of mRNA expression of 6 ISGs, see Methods) (**I**) and *IFNB1* mRNA expression (**J**) in MTPAP_KO cells compared to controls measured by qPCR. Mean ± SEM; n=7 experiments; each colour corresponds to KO_MTPAP cells generated using a single guide RNA targeting MTPAP; ** p<0.01 in Kruskall-Wallis with Dunn’s post-hoc test.

**Table 1:** Summary of molecular and clinical data in patients with mutations in *MTPAP* included in this study.

| Patient | Patient ID | *MTPAP* variant(s) | Effect on protein/activity | Phenotype | Reference |
| --- | --- | --- | --- | --- | --- |
| P1 | AGS0432.1 | c.1153A>T (p.Ile385Phe) (HOM) | Unknown | Lethal encephalopathy | Van Eyck et al., 2020 |
| P2 | AGS3853.1 | c.245G>A (p.Cys82Tyr) (HOM) | Reduced protein level | Ataxia, microcephaly | This study |
| P3 | AGS0664.1 | c.1283T>C (p.Ile428Thr); c.1567C>T (p.Arg523Trp) | Reduced protein level, impaired activity | Perinatal encephalopathy (AGS-like) | Van Eyck et al., 2020 |
| P4 | AGS0664.4 | c.1283T>C (p.Ile428Thr); c.1567C>T (p.Arg523Trp) | Reduced protein level, impaired activity | Perinatal encephalopathy (AGS-like) | Van Eyck et al., 2020 |
| P5 | AGS4181.1 | c.1432A>G (p.Asn478Asp) (HOM) | Reduced protein level, no activity | Complex spastic ataxia, optic atrophy | Crosby et al., 2010; Wilson et al., 2014 |

AGS = Aicardi-Goutières syndrome ; HOM = Homozygous

To elucidate the link between loss of MTPAP and IFN signalling, we deleted *MTPAP* by CRISPR/Cas9 using three different guides in the BJ-5ta fibroblast cell line (Figure 1B). Similar to the phenotype observed in patient fibroblasts, knocking-out (KO) *MTPAP* strongly reduced the levels of COX2, an mtDNA-encoded subunit of complex IV, and NDUFB8, a nuclear-encoded participant of complex I, which includes several mtDNA-encoded subunits (Figure 1A-1B). Other subunits of OXPHOS were less severely affected, except in cells from P5 (Supp Figure 1B-C).

Upon evaluation of the mitochondrial network in MTPAP_KO cells and patient fibroblasts using Fiji *MiNA* plugin (Valente et al., 2017), we did not observe any significant change in mitochondrial area, measured as *footprint,* compared to controls (Figure 1C-D). Consistently, mtDNA content was not reproducibly affected, except for an increase in fibroblasts from P4 (Supp Figure 1D-E). Summarising these data, mitochondrial mass and DNA content were minimally affected by the loss of MTPAP.

Transmission electron microscopy of patient fibroblasts with *MTPAP* mutations suggested abnormal cristae, folds of the inner membrane where respiratory chain complexes and ATP synthase are found (Genin et al., 2015), compared to controls (Supp Figure 1F). In line with this feature of mitochondrial damage, mitochondrial membrane potential, measured using tetramethyl rhodamine ethyl ester (TMRE), was reduced in both KO cells and patient primary fibroblasts compared to controls (Figure 1E-1F), indicating impaired mitochondrial function. We further characterised mitochondrial respiration using the Seahorse *Mito Stress* assay in MTPAP_KO cells, observing a significant reduction of ATP production and basal respiration compared to controls (Figure 1G-H). Similar results were recorded in fibroblasts from P2 (Supp Figure 1G-H). In previous reports, while fibroblasts from P3 and P4 showed only a moderate alteration in respiratory capacity (Van Eyck et al., 2020), fibroblasts from P5 also showed a loss of complex I and complex IV activity (Wilson et al., 2014).

Strikingly, we observed a marked upregulation of ISGs and *IFNB1* mRNA, measured by qPCR, in MTPAP_KO BJ-5ta cells compared to controls (Figure 1I-J), consistent with our findings in the blood of P1 and P2.

Summarising, the above data indicate that MTPAP dysfunction is associated with loss of some OXPHOS subunits and impaired mitochondrial respiration. Additionally, we recorded an upregulation of ISGs and *IFNB1*, suggesting a link between MTPAP dysfunction and type I IFN signalling upregulation.

### Mitochondrial nucleic acid sensing involves the cytosolic RNA sensor MDA5 and its adaptor MAVS

Mitochondrial nucleic acid depletion with dideoxycytidine (ddC) (Brown & Clayton, 2002) in MTPAP_KO cells abolished ISGs and *IFNB1* upregulation (Figure 2A-B, Supp Figure 2A), indicating that the induction of type I IFN signalling observed in the absence of MTPAP is due to mitochondrial DNA and/or RNA sensing.

**Figure 2:**
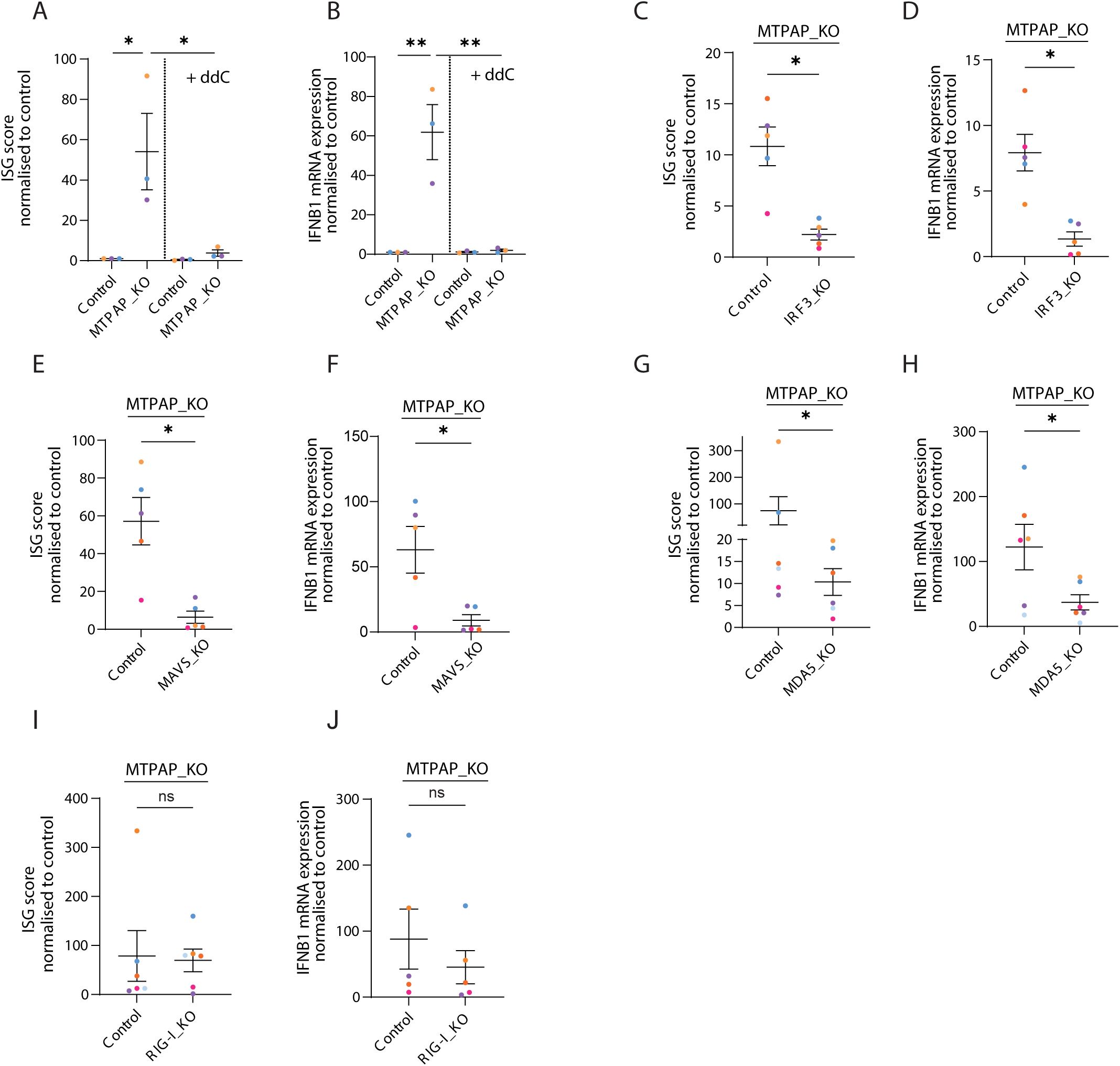
Mitochondrial nucleic acid sensing involves the cytosolic RNA sensor MDA5 and its adaptor MAVS. **A-B:** ISG score (**A**) and *IFNB1* mRNA (**B**) in control and MTPAP_KO (sg#2) cells treated for 10 days with ddC. Mean ± SEM; n=4, each colour is a different experiment; * indicates p<0.05, ** p< 0.01, 2-way ANOVA with Holm-Sidak multiple comparison test. **C-D:** ISG score (**C**) and *IFNB1* mRNA levels (**D**) in BJ-5ta cells double KO IRF3/MTPAP compared to controls. **E-F**: ISG score (**E**) and *IFNB1* mRNA levels (**F**) in MAVS/MTPAP double KO cells compared to controls. **G-H**: ISG score (**G**) and *IFNB1* mRNA levels (**H**) in MDA5/MTPAP double KO cells compared to controls. **I-J**: ISG score (**I**) and *IFNB1* mRNA levels (**J**) in RIG-I/MTPAP double KO cells compared to controls. **C-J:** Mean ± SEM; n≥4, each colour is a different experiment, KO_MTPAP data are represented by the average of sg#1, sg#2 and sg#3 data; ns indicates non significance, * indicates p<0.05 in Wilcoxon test.

In order to determine the pathway involved in mitochondrial nucleic acid detection upon loss of MTPAP, we deleted molecules known to be involved in cytosolic nucleic acid sensing and IFN signalling in MTPAP_KO cells. As expected, KO of IRF3, a transcription factor responsible for *IFNB1* induction (De Lendonck et al., 2014), abolished *IFNB1* expression and ISG upregulation in MTPAP_KO cells (Supp Figure 2B; Figure 2C-D). While deletion of the DNA sensor cGAS and its adaptor STING abolished IFN responses to transfected DNA (Supp Figure 2C-D-E), it did not reduce IFN signalling triggered by MTPAP deletion (Supp Figure 2F-G-H-I), indicating that DNA sensing was not involved. Conversely, deletion of MAVS, the essential adaptor protein involved in cytosolic RNA detection through the dsRNA sensors RIG-I and MDA5, was associated with a marked reduction of *IFNB1* and ISG induction following MTPAP loss (Supp Figure 2J-2K; Figure 2E-F). To confirm the involvement of RNA sensing, we deleted MDA5 and RIG-I in MTPAP_KO cells (Supp Figure 2L, 2N). Only the KO of MDA5 led to a reduction in ISGs and *IFNB1* expression (Figure 2G-J). As controls, cytosolic RNA sensing due to dsRNA poly(I:C) transfection and Sendai virus exposure was strongly affected by loss of MAVS, MDA5 and RIG-I (Supp Figure 2M, 2O). Summarising, the above data show that, in the absence of MTPAP, mitochondrial nucleic acids trigger an IRF3-mediated innate immune response involving MAVS and MDA5, suggesting the sensing of dsRNA of mitochondrial origin.

### Cytosolic leakage of mt-dsRNA triggers innate immune signalling

Given the above results, and a previous report describing an accumulation of mt-dsRNA in the absence of MTPAP in Drosophila (Pajak et al., 2019), we assessed the presence of dsRNA in control and MTPAP_KO cells. Immunostaining using J2 antibody, which detects dsRNA molecules longer than 40 nucleotides (Schönborn et al., 1991; Weber et al., 2006), revealed a marked accumulation of dsRNA in mitochondria in the absence of MTPAP, whereas very little signal was detected in control cells (Figure 3A-B). Of note, the dsRNA foci observed in the absence of MTPAP were larger than the few detected in control cells (Figure 3C). In patient cells, the dsRNA signal in mitochondria was also increased (Supp Figure 3A-B), with a larger proportion of cells with high dsRNA signal compared to controls (Supp Figure 3C). Interestingly, these dsRNA structures disappeared when transcription was inhibited by actinomycin D treatment (Supp Figure 3D-E), suggesting that they originate from recently transcribed RNA. Strikingly, similar to IFN signalling (Fig 2A-B), the enlarged dsRNA foci in MTPAP_KO cells disappeared upon depletion of mitochondrial nucleic acids with ddC (Supp Figure 3F). We also verified the specificity of the J2 signal using dsRNA-specific RNase III (Supp Figure 3F). These data suggest that the accumulated dsRNA seen in the context of MTPAP deletion are generated through mtRNA transcription.

**Figure 3:**
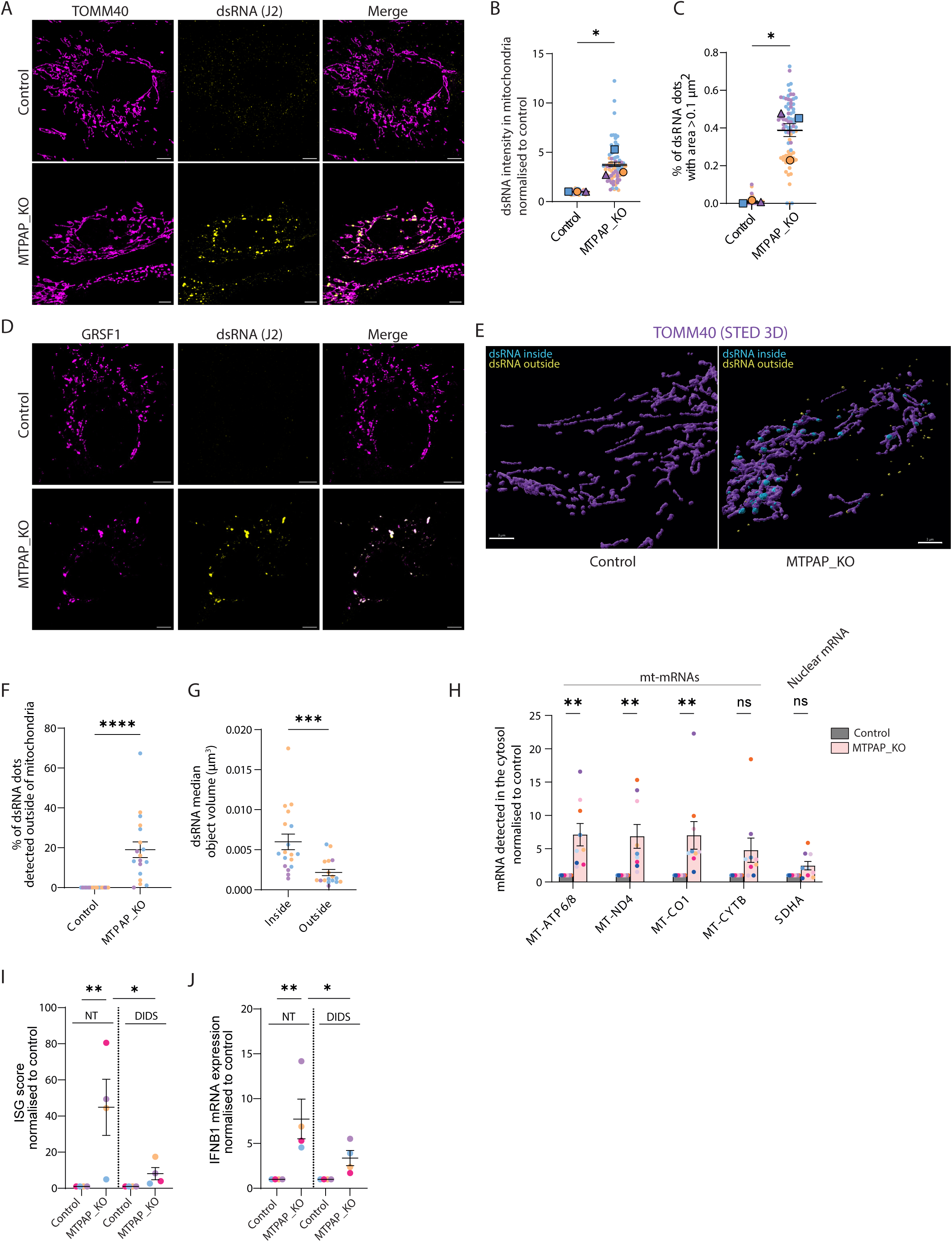
mt-dsRNAs are released into the cytosol via mitochondrial pores. **A:** Representative confocal microscopy image of immunostaining with antibody to mitochondrial protein TOMM40 and J2 antibody to dsRNA in control and MTPAP_KO (sg#1) cells. Scale bar: 5 µm. **B:** Quantification of average pixel intensity of J2 dsRNA signal in mitochondria in control vs MTPAP_KO cells (average of sg#1, #2 and #3). Superplot, mean ± SEM; n=3 experiments; each borderless point represents the measure of a cell; each colour is a different experiment; * indicates p<0.05 in paired t-test performed on the mean value for each experiment. **C:** Percentage of dsRNA dots in mitochondria with an area over 1 µm^2^ in control cells vs MTPAP_KO cells (average of sg#1, #2 and #3). Superplot, mean ± SEM; n=3 experiments; each borderless point represents the measure of a cell; each colour is a different experiment; * indicates p<0.05 in paired t-test performed on the mean value for each experiment. **D:** Representative confocal microscopy image of immunostaining with antibody to mitochondrial RNA granule protein GRSF1 and dsRNA (J2) in control and MTPAP_KO (sg#2) BJ-5ta cells. Scale bar: 5 µm. **E:** Representative image of the 3D volume reconstruction of STED images after immunostaining with antibody against mitochondrial protein TOMM40 and dsRNA (J2) in control and MTPAP_KO cells (sg#2). Yellow dots correspond to dsRNA particles detected outside of the mitochondrial network. Scale bar: 3 µm. **F:** Percentage of dsRNA dots detected outside of the mitochondrial network in control vs MTPAP_KO cells (sg#2). Mean ± SEM; n=3 experiments; each point represents the measure of a cell; each colour is a different experiment; **** indicates p<0.0001 in Mann-Whitney test. **G:** Quantification of the median volume of dsRNA particles detected inside or outside of the mitochondrial network in MTPAP_KO cells (sg#2). Mean ± SEM; n=3 experiments; each point represents the measure of a cell; each colour is a different experiment; *** indicates p<0.001 in Mann-Whitney test. **H:** Quantification of mtRNA levels measured by qPCR in cytosolic fractions of control and MTPAP_KO cells (sg#2). Mean ± SEM; n=8 experiments; each colour is a different experiment; ns indicates non significance, ** p< 0.01 in 2-way ANOVA with Holm-Sidak multiple comparison test. **I-J:** ISG score (**I**) and *IFNB1* mRNA expression (**J**) measured by qPCR in MTPAP_KO cells (average of sg#1, sg#2 and sg#3 data for each experiment) treated for 24h with 300 µM DIDS compared to controls. Mean ± SEM; n=4 experiments, each colour is a different experiment; * indicates p<0.05, ** p<0.01, two-way ANOVA with Holm-Sidak multiple comparison test.

Consistently, in MTPAP_KO cells, mt-dsRNA colocalized with mtRNA granule protein GRSF1 (Jourdain et al., 2014), which appeared to re-localise within mitochondria together with dsRNA foci (Figure 3D). Interestingly, mt-dsRNA also colocalized with the mitochondrial degradosome protein PNPase (Supp Figure 3G), involved in mt-dsRNA elimination (Dhir et al., 2018). To corroborate the results of our confocal microscopy analyses, we performed super-resolution microscopy (STED 3D) on immunostained dsRNA and mitochondria. 3D volume analysis using *Imaris* indicated that the particles of dsRNA were not only bigger, but also less round in MTPAP_KO cells compared to control cells (Supp Figure 3H-I), suggesting structures distinct from homeostatic MRGs.

Notably, 3D reconstruction of the mitochondrial network revealed the presence of dsRNA foci outside of mitochondria in the absence of MTPAP (Yellow dots on Figure 3E). More precisely, in MTPAP_KO cells, 20 % of dsRNA particles localized outside of the mitochondrial network (Figure 3F), and were significantly smaller than the ones retained within mitochondria (Figure 3G). To confirm the mitochondrial origin of this cytosolic dsRNA, we employed cellular fractionation using the mild detergent digitonin, differential centrifugation, and then qPCR on RNA from the cytosolic fraction (Bryant et al., 2022; Hootman et al., 2023) (Supp Figure 3J-K). We detected higher levels of several mtRNAs, in particular *MT-ATP6/8*, *MT-ND4*, *MT-CO1*, in the cytosol of MTPAP_KO cells compared to control cells (Figure 3H), consistent with the release of mt-dsRNA into the cytosol. Further, treatment with the VDAC1 inhibitor DIDS, implicated in cytosolic release of mtDNA (Kim et al., 2019) and mtRNA (Krieger et al., 2024), reduced the upregulation of type I IFN signalling due to loss of MTPAP (Figure 3I-J). Summarising, these data indicate that the absence of MTPAP results in the accumulation, and cytosolic leakage, of mt-dsRNA through VDAC1 channels, leading to IFN signalling.

### MTPAP loss results in a reduction of mt-mRNA levels in parallel with an accumulation of non-coding mtRNA

While a previous study recorded a loss of mt-mRNA upon dysfunction of MTPAP (Wilson et al., 2014), we observed an accumulation of mt-dsRNA in the absence of MTPAP, suggesting the need for a broad survey of mtRNA species. We measured processed and unprocessed mitochondrial transcripts using NanoString technology and a ‘Mitostring’ custom set of probes (Wolf and Mootha, 2014; Foged et al., 2025). Notably, this approach allows for the concomitant quantification of all mt-mRNA, junction mtRNA (mtRNA sequences between two adjacent coding genes) including mt-tRNAs before their excision, noncoding mtRNA originating from mtDNA heavy or light strand transcripts, and selected nuclear-encoded mRNA (Figure 4A). As expected, we observed reduced levels of several mt-mRNAs, including *MT-ATP6*, *MT-CO1, MT-CO2* and *MT-CYTB* in both MTPAP_KO cells and patient fibroblasts compared to controls (Figure 4B-C). Contrastingly, levels of *MT-ND1* and *MT-ND2* mt-mRNAs were elevated in MTPAP_KO cells, indicating variable effects of MTPAP dysfunction on mt-mRNA stability (Figure 4B-C). Levels of *MT-ND6*, reported as non-polyadenylated (Ruzzenente et al., 2011), were unaffected by the loss of MTPAP in patient cells, and increased in MTPAP_KO cells (Figure 4B-C). Further, the abundance of junction mtRNA and mt-tRNA remained largely unchanged by MTPAP dysfunction, suggesting that mtRNA transcription was not modified (Supp Figure 4A-B). Of particular note, we observed a striking increase in the levels of one non-coding mtRNA sequence, detected with the probe for light strand reverse complementary RNA of *MT-CO1* (rc*_MT-CO1*), in both MTPAP_KO cells and patient fibroblasts (Figure 4D-E). This was also evident with the mtRNA junction probe rc*_MT-CO1_TRNS1* (Supp Figure 4A-B). Nuclear-encoded mRNA of proteins involved in mtRNA processing were not significantly changed, except for *MTPAP* in patient cells while it was unchanged in MTPAP_KO as CRISPR/Cas9 edition does not necessarily affect mRNA levels (Zhu et al., 2024) (Supp Figure 4C-D). Of note, the abundance of mt-rRNA, which were not included in the MitoString probe set, was measured by qPCR and found to be unaffected upon loss of MTPAP (Supp Figure 4E-F).

**Figure 4:**
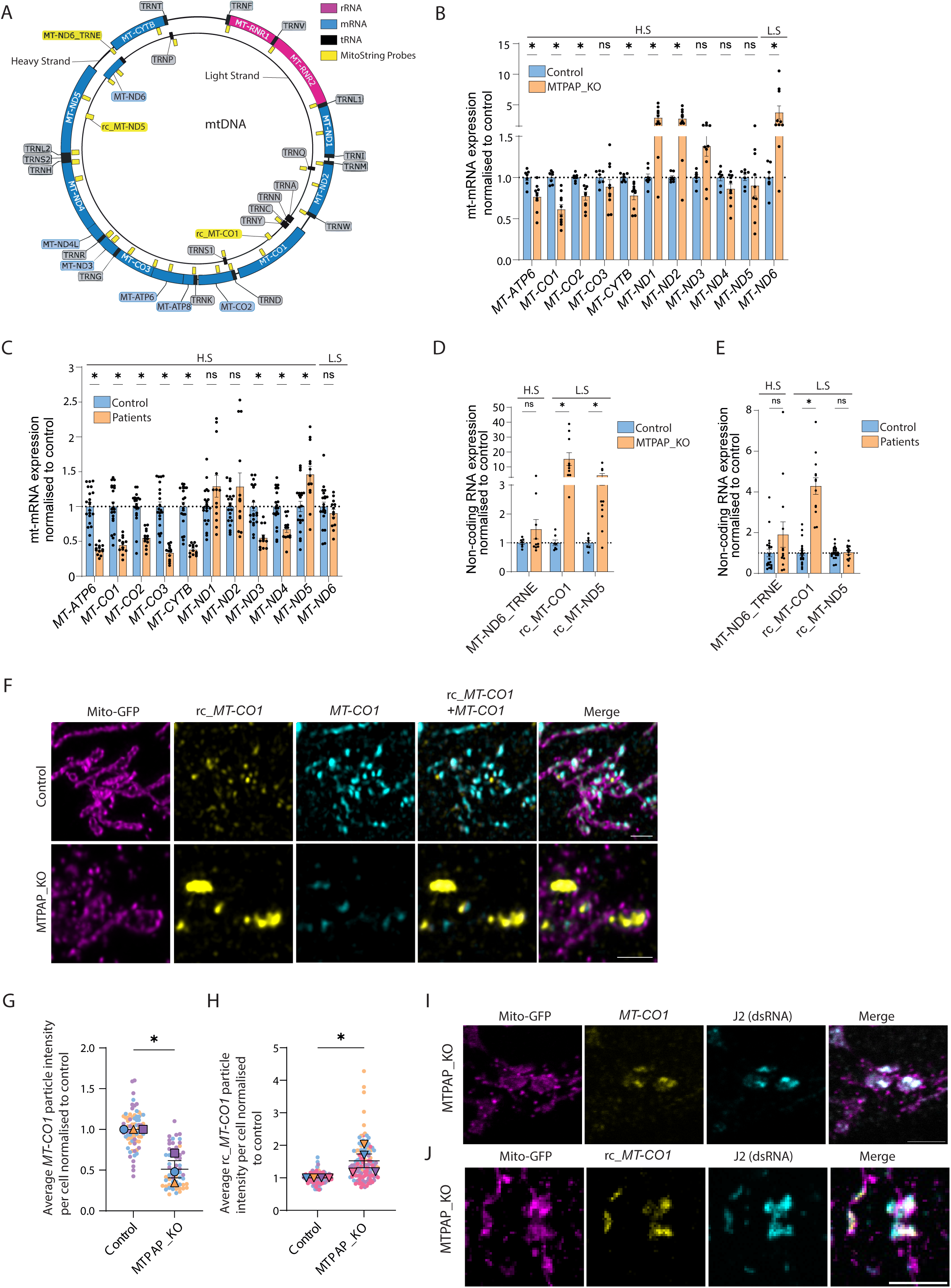
MTPAP loss causes reduced mt-mRNA levels in parallel with accumulation of noncoding mtRNA. **A:** Mapping of transcripts targeted by MitoString probes on a schematic representation of mtDNA. **B-C:** Relative levels of mt-mRNA in MTPAP_KO cells (sg#1, #2, #3 each collected 3 times) (**A**) and fibroblasts from patients with *MTPAP* mutations (P2, P3, P4, P5 each collected 3 times) (**B**) compared to respective controls and quantified using MitoString. H.S.: heavy strand, L.S.: light strand. Mean ± SEM; n=3 experiments; each point corresponds to a different sample, ns indicates non significance, * p< 0.05 in multiple Mann-Whitney with Holm-Sidak comparison tests. **D-E:** Relative levels of non-coding mtRNA in MTPAP_KO cells (sg#1, #2, #3 each collected 3 times) (**D**) and of patients with *MTPAP* mutations (P2, P3, P4, P5 each collected 3 times) assessed using MitoString compared to controls. H.S. : heavy strand, L.S. : light strand. Mean ± SEM; n=3 experiments; each point corresponds to a different sample, * indicates p< 0.05 in multiple Mann-Whitney with Holm-Sidak comparison tests. **F:** Representative confocal microscopy images of Mito-GFP mitochondrial marker and sm-FISH probes *MT-CO1*-Cy5 and rc_*MT-CO1*-Cy3 in control and MTPAP_KO cells (sg#1) from 3 experiments. Scale bar: 5µm. **G-H:** Quantification of the average fluorescence intensity signal per cell of single particles detected with sm-FISH probe *MT-CO1*-Cy3 (**G**) and rc_*MT-CO1*-Cy3 (**H**) in control and MTPAP_KO cells (average of sg#1 and sg#2). Superplot, mean ± SEM; n≥3 experiments; each borderless point represents the measure of a cell; each colour is a different experiment; * indicates p<0.05 in paired t-test performed on the mean value for each experiment. **I:** Representative confocal microscopy images of Mito-GFP mitochondrial marker, smFISH probes *MT-CO1*-Cy3, and immunostaining of dsRNA with J2 antibody in MTPAP_KO cells from 3 experiments. Scale bar: 5 µm. **J:** Representative confocal microscopy images of Mito-GFP mitochondrial marker, smFISH probes rc_*MT-CO1*-Cy3, and immunostaining of dsRNA with J2 antibody in MTPAP_KO cells from 3 experiments. Scale bar: 5 µm.

Accumulation of rc_*MT-CO1*, in the presence of reduced but not abolished levels of its complementary RNA species *MT-CO1*, suggested a potential contribution to the formation of immunostimulatory mt-dsRNA in MTPAP_KO cells. Indeed, according to absolute RNA molecule quantification by MitoString, there was approximately one rc_*MT-CO1* RNA molecule for every 500 *MT-CO1* molecules under control conditions. In contrast, in MTPAP_KO cells and in patient fibroblasts, the loss of *MT-CO1* and the accumulation of rc_*MT-CO1* raised this ratio to about 1:22 and 1:46 respectively, greatly increasing the likelihood of complementary strand hybridization. Using single molecule FISH (sm-FISH) (Tsanov et al, 2016; Safieddine et al., 2022), we were able to detect both *MT-CO1* RNA and rc_*MT-CO1* in mitochondria (Figure 4F). Consistent with above results, in the absence of MTPAP, the overall sm-FISH signal of *MT-CO1* was reduced (Figure 4F-G), while that of rc_*MT-CO1* was increased (Figure 4F, H). Further, the remaining *MT-CO1* and accumulated rc_*MT-CO1* RNAs partially overlapped in MTPAP_KO cells, supporting their contribution to the formation of mt-dsRNA (Figure 4F). Importantly, both *MT-CO1* and rc_*MT-CO1* sm-FISH signals were present within dsRNA structures, stained with J2 antibody and accumulating in MTPAP defective cells (Figure 4I). Interestingly, *MT-CO1* was among the mtRNA observed to leak into the cytosol in MTPAP_KO cells (Fig 3H). These data suggest that, upon loss of MTPAP, *MT_CO1* RNA and rc_*MT-CO1* likely contribute to the formation of mt-dsRNA foci, followed by their release into the cytosol and subsequent sensing by the innate immune machinery.

## Discussion

In this study, we show that loss of MTPAP activity triggers limited changes in mitochondrial respiratory chain protein levels while inducing significant mitochondrial damage, reduced mitochondrial membrane potential and impaired mitochondrial respiration. In addition, upon loss of MTPAP, we recorded an upregulation of type I IFN signalling that was induced by the release of mtRNA into the cytosol involving VDAC1 channels, and detection through the MDA5/MAVS sensing pathway. We observed reduced levels of some mt-mRNAs, consistent with MTPAP dysfunction leading to loss of poly-A tails on mt-mRNAs and their subsequent degradation. However, we also observed a marked accumulation of light-strand derived noncoding mtRNA, most notably the complementary strand of *MT-CO1*, colocalizing with large dsRNA structures involving MRG components. Thus, our results reveal an unexpected role for MTPAP in mtRNA turnover, preventing the accumulation of immunostimulatory mtRNA in mitochondria and their release into the cytosol. Importantly, in patients with mutations in *MTPAP,* we recorded type I IFN signalling upregulation which might directly contribute to pathogenesis.

MTPAP is necessary for maturation of mt-mRNAs through polyadenylation, and for monoadenylating mt-tRNA (Fiedler et al., 2015). In particular, loss of MTPAP function leads to destabilisation and reduced levels of some mt-mRNAs such as *MT-CO1* and *MT-CO3* (Wilson et al., 2014), consistent with our findings in both patient fibroblasts and MTPAP_KO cells. Using distinct detection techniques, two large scale studies of mtRNA metabolism reported contrasting effects of MTPAP depletion, i.e., decreased (Wolf and Mootha, 2014; Foged et al., 2025), or increased (Kim et al., 2024) mtRNA levels. Interestingly, our broad survey of mtRNA with MitoString sheds light on this apparent discrepancy, revealing that mt-mRNA transcripts were differentially affected by the loss of MTPAP. Thus, levels of *MT-ND1* and *MT-ND2* were increased, as in Wilson et al. (Wilson et al., 2014), while *MT-ND5* and *MT-ND6* were mainly unaffected upon MTPAP dysfunction. In contrast, most other mt-mRNAs - not analysed in the study of Kim et al. (Kim et al., 2024) - were downregulated. *MT-ND6*, the only mt-mRNA encoded on the light strand, is non-polyadenylated, consistent with a stability maintained upon MTPAP loss. Polyadenylation of *MT-ND1* and *MT-ND2* mRNA has not been described as distinct from other mt-mRNAs, while *MT-ND5* poly-A tails might be slightly shorter than other transcripts according to a recent report (McShane et al. 2024), questioning the relationship between polyadenylation and stability. Of note, similar amounts of junction mtRNAs suggest that mitochondrial transcription and primary transcripts levels are unaffected by loss of MTPAP function, and do not explain changes in mtRNA abundance. Notably, according to a recent study (McShane et al., 2024), *MT-ND1* and *MT-ND2* are significantly less abundant and shorter lived than other mt-mRNAs, suggesting a faster turnover mediated by a higher rate of degradation. Thus, MTPAP and polyadenylation may have a role in promoting the degradation, rather than the stabilisation, of some mtRNAs, consistent with their accumulation when MTPAP activity is lost.

Interestingly, it has been reported that MTPAP adds long poly-A tails to defective mt-tRNAs, marking them for degradation by the degradosome (Pearce et al., 2017; Tompuu et al., 2018). Thus poly-A tails may constitute a signal for degradation, with MTPAP dysfunction leading to impaired degradation and mtRNA accumulation. Determinants of the stability- versus degradation- promoting roles of poly-A tails remain to be elucidated. Strikingly, we observed accumulation of noncoding transcripts in MTPAP defective cells, such as the complementary strand of *MT-CO1*, which likely arise from defective degradation of the light strand derived transcript by the degradosome. Of note, rc-*MT-CO1* accumulation was also detected in mitochondrial protein gene silencing screens including MTPAP (*Wolf & Mootha, 2014*; Foged et al., 2025). Therefore, MTPAP may be involved in adding a poly-A “degradation signal” to such potentially detrimental noncoding mtRNA species, so as to ensure their very low level, similarly to the uridylation-mediated decay of mtRNA in *Trypanosoma Brucei* (Ryan & Read, 2005).

Accumulated noncoding mtRNAs in MTPAP defective cells have the potential to hybridize with residual complementary mt-mRNA to form immunostimulatory mt-dsRNA. Indeed, we observed a striking accumulation of dsRNA foci of mitochondrial origin, with evidence of the contribution of rc-*MT-CO1* and *MT-CO1.* This is in agreement with the description of dsRNA accumulation in an MTPAP null *Drosophila* model (Pajak et al., 2019). Mt-dsRNA formed large granules colocalizing with GRSF1 and PNPase, involved in mtRNA processing in MRGs (Szczesny et al. 2010; Jourdain et al. 2014). However, these granules did not result from transcriptional inhibition as described recently (Hansen et al., 2025), since levels of the primary transcripts were unaffected, and dsRNA foci disappeared when total transcription was inhibited by actinomycin D, or when the template for mtRNA, i.e. mtDNA, was depleted. Conversely, the mt-dsRNA granules we observed could be similar to mitochondrial stress bodies (MSB), reported to sequester mtRNA as a response to proteostatic stress when SUV3, a component of the degradosome, is lost (Begeman et al., 2025). In MTPAP_KO cells, *SUPV3L1* mRNA levels were unaffected, but the accumulation of mt-dsRNA suggests that other determinants of degradosome activity are impaired, because of MTPAP dysfunction.

Importantly, we show that mt-dsRNAs can reach the cytosol, where they can be sensed by the MDA5/MAVS pathway and trigger type I IFN signalling. This possibility is consistent with the findings of Dhir et al. and Idiiatullina et al., describing defective mtRNA degradation due to PNPase and REXO2 deficiencies respectively, leading to mt-dsRNA accumulation, mt-dsRNA cytosolic release, and induction of a mtRNA-dependent type I IFN response (Dhir et al., 2018; Idiiatullina et al., 2024). Although the process of mt-dsRNA leakage upon loss of MTPAP remains largely unclear, our data indicate a contribution for VDAC1 channels, as previously reported (Krieger et al., 2024).

Since 2018, multiple studies have described mtRNA accumulation and release from mitochondria in various contexts, including a disturbance of mtRNA at distinct processing steps (Foged et al., 2025; Yoon et al., 2025), inhibition of the ATP synthase (Kim et al., 2022), and deficiency of fumarate hydratase (Hootman et al., 2023) or caspases (Killarney et al., 2023). Thus, stressors with no clear link to mtRNA processing *per se* can induce an increase in dsRNA levels in mitochondria, and their subsequent cytosolic release. These observations raise the possibility that dsRNA accumulation constitutes a ‘generic’ indicator of mitochondrial stress, with immunostimulatory potential. While in these cases, the molecular basis for dsRNA accrual is unclear, we show that loss of MTPAP function has a specific effect on some noncoding mtRNAs, which may be polyadenylated prior to their processing.

Importantly, our study describes MTPAP deficiency to be associated with enhanced type I IFN signalling, as demonstrated in patients carrying mutations in *PNPT1* (Dhir et al 2018). Thus, although very few individuals harbouring biallelic mutations in *MTPAP* have been reported to date (Ravanbod et al., 2025), this study adds to the list of mitochondrial defects leading to IFN signalling upregulation in humans (Lepelley et al., 2021a; Crow and Casanova, 2024). Given the challenges in diagnosing and treating mitochondrial diseases (Khan et al., 2015; Zheng et al., 2024), recognition of an inflammatory component in certain mitochondrial cytopathies may help improve diagnosis and patient care, raising the question of the potential efficacy of ‘anti-interferon’ therapies in such cases.

## MATERIAL & METHODS

### Samples obtained from patients

The study was approved by the Comité de Protection des Personnes (ID-RCB/EUDRACT: 2014-A01017-40; revalidated in 2025).

### Clinical data for P2

We report on an unpublished male patient from Sweden, the first child of healthy consanguineous parents of Asian origin. The patient presented with global psychomotor delay, first noted at 7 months of age, and developed progressive microcephaly and ataxia during infancy. Clinical progression was noted during the first year, followed by a relatively stable neurological course thereafter.

At two years of age, brain MRI demonstrated periventricular leukomalacia accompanied by multiple foci of increased T2 signal intensity within the supratentorial and subcortical white matter, without evidence of a generalized leukoencephalopathy. Brain CT showed no intracranial calcifications, thereby making Aicardi-Goutières syndrome unlikely. Ophthalmological examination excluded optic atrophy; however, the child exhibited bilateral severe sensorineural hearing loss. Neurological examination revealed generalized muscle weakness, absent tendon reflexes in the lower limbs, and signs consistent with peripheral neuropathy confirmed by neurography. Muscle biopsy showed normal histological architecture and respiratory chain enzyme activity.

Genetic testing identified a homozygous missense variant in the *MTPAP* gene (NM_018109.4), c.245G>A leading to a p.Cys82Tyr substitution (not seen on GnomAD v4.1), classified as a variant of uncertain significance (VUS). Both parents were confirmed as heterozygous carriers.

### IFN score in patient blood

Whole blood was collected into PAXgene tubes (Qiagen), and total RNA was extracted using a PreAnalytix RNA isolation kit. IFN scores were generated in one of two ways as previously described. For P1, TaqMan probes and qPCR were used to measure the mRNA expression of six ISGs (IFI27, IFI44L, IFIT1, ISG15, RSAD2, and SIGLEC1) normalized to the expression level of HPRT1 and 18S rRNA (Rice et al., 2013). The median fold change of the ISGs is compared with the median of 29 healthy controls to create an IFN score for each individual, with an abnormal score being defined as >2.466. For P2, NanoString technology was employed. Analysis of 24 genes and 3 housekeeping genes (probes of interest [*n* = 24]: *IFI27*, *IFI44L*, *IFIT1*, *ISG15*, *RSAD2*, *SIGLEC1*, *CMPK2*, *DDX60*, *EPSTI1*, *FBXO39*, *HERC5*, *HES4*, *IFI44*, *IFI6*, *IFIH1*, *IRF7*, *LAMP3*, *LY6E*, *MX1*, *NRIR*, *OAS1*, *OASL*, *OTOF*, and *SPATS2L*; reference probes [*n* = 3]: *NRDC*, *OTUD5*, and *TUBB*) was conducted using the NanoString customer designed CodeSet according to the manufacturer’s recommendations (NanoString Technologies). Agilent Tapestation was used to assess the quality of the RNA. 100 ng total RNA was loaded for each sample. Data were processed with nSolver software (NanoString Technologies). Data were normalized to the internal positive and negative calibrators, the three reference probes, and the control samples. The median of the 24 probes for each of the 27 healthy control samples was calculated. The mean NanoString score of the 27 healthy controls +2 SD of the mean was calculated. Scores above this value (2.758) were designated as positive.

### Cell culture

Primary fibroblasts (from three healthy donors (controls), and patients 2, 3, 4 and 5) and BJ-5ta fibroblasts (catalog no. CRL-4001; ATCC) were grown in DMEM supplemented with 10% (vol/vol) fetal bovine serum. For primary fibroblasts, 1% penicillin-streptomycin and 0.25 µg/ml amphotericin B (GIBCO) were added to the media. All cell lines were maintained at 37°C in 5% CO_2_.

### VDAC1 inhibition

DIDS (4,4’-diisothiocyanostilbene-2,2’-disulfonic acid; 309795; Sigma-Aldrich), a stilbene disulfonate anion channel inhibitor that binds VDAC1 and prevents its oligomerization and conductance, stock (100 mM) was prepared in DMSO and stored at -20°C protected from light. VDAC1 activity was blocked using 300 µM DIDS for 24h.

### Cell transcription arrest

Cell transcription was inhibited using actinomycin D (ActD) at 2.5 µg/mL during a 24 h time course to monitor dsRNA disappearance kinetics. Cells were either lysed or fixed at 2, 8, 16, and 24 h post ActD treatment initiation and compared to non-treated controls.

### Generation of knock-out cells by CRISPR/Cas9

Single guide RNAs targeting *MTPAP, IRF3, cGAS, STING, MAVS, MDA5, RIG-I* (oligonucleotides sequences are in **Supp Table 1**) were previously described (Johnson et al.,2017; Dunker et al., 2021) or designed using the CRISPOR tool (crispor.tefor.net) and cloned into lentiviral constructs lentiCRISPRv2 (a gift from Feng Zhang, Addgene plasmid # 52961) (Sanjana et al., 2014) or lentiCRISPRv2-hygro (a gift from Brett Stringer, Addgene plasmid # 98291) (Stringer et al., 2019), following the protocol provided on Addgene.

Lentiviral vectors carrying these constructs were produced by calcium phosphate transfection of 293FT cells in combination with packaging vectors psPAX2, (psPAX2 was a gift from Didier Trono, Addgene plasmid # 12260), and envelope pCMV-VSV-G (pCMV-VSV-G was a gift from Bob Weinberg, Addgene plasmid # 8454) (Stewart et al., 2003). Medium of 70% confluent 293FT in 75-cm^2^ flasks was changed 2 h before transfection. Calcium phosphate precipitates were prepared by mixing 12.5 µg lentiviral vectors with 12.5 µg psPAX2 and 5 µg pCMV-VSV-G in water for a final volume of 875 µL. 125 µL 2 M CaCl_2_ and 1 mL HBS 2X (50 mM Hepes, 10 mM KCl, 280 mM NaCl, and 1.5 mM Na_2_HPO_4_, pH 7.05) were sequentially added dropwise in slowly vortexed solution. Solutions were incubated at room temperature for 20 min and mixed gently with 293FT supernatant. Medium was replaced by 7 mL culture medium 24 h later. After 24 more hours, supernatants were collected, centrifuged at 1,700 rpm for 5 min and 0.45-µm filtered. 250 000 BJ-5ta cells were transduced with 1 mL lentiviral vector supernatant, 8 µg/mL polybrene (Millipore), and 10 mM Hepes (Invitrogen) in 6-well plates and medium replaced 24 h later. 2 days after transduction, transduced cells were selected with 10 µg/mL puromycin (Sigma-Aldrich) or 200 µg/mL hygromycin B (Invivogen) Cells were collected for analysis starting from 14 days after transduction, and KO efficiency verified by western blotting and functional assays.

For sm-FISH, we used a BJ-5ta cell line stably expressing the Mito-GFP construct pMXs-3XHA-EGFP-OMP25 (a gift from David Sabatini, Addgene plasmid # 83356) (Chen et al., 2016), after transduction as described above and flow cytometry sorting to select for a homogenous EGFP expression level.

### Western blot analysis

For whole-cell lysate analysis, proteins were extracted using lysis buffer (radioimmunoprecipitation assay, 1% protease inhibitor, and 1% phosphatase inhibitor). Bolt LDS Sample Buffer (4×; Novex Life Technologies) and Bolt Sample Reducing agent (10×; Novex Life Technologies) were added to protein lysates; samples were resolved on 4–12% Bis-Tris Plus gels (Invitrogen) and then transferred onto nitrocellulose membranes for 7 min at 20 V using the iBlot 3 Dry Blotting System (ThermoFisher). Membranes were blocked with 5% non-fat milk in Tris-buffered saline (TBS) and primary antibodies incubated overnight at 4°C in 5% non-fat milk in TBS buffer supplemented with 0.1% Tween. A list of antibodies used in this study is supplied in **Supp Table 2**. Membranes were washed and incubated with appropriate fluorescent anti-mouse or anti-rabbit secondary antibodies (secondary antibody list in **Supp Table 3**) for 45 min at room temperature (LI-COR System). Signal was detected using the Odyssey CLx System (LI-COR). Signal quantifications were performed using Image Studio (LI-COR).

### RT-qPCR quantification of gene expression

Total RNA was extracted using the RNAqueous-Micro Kit (Ambion), and reverse transcription was performed with the High-Capacity cDNA Reverse Transcription Kit (Applied Biosystems). Levels of cDNA were quantified by RT-qPCR on 25 ng/µL cDNA using TaqMan Gene Expression Assays (Applied Biosystems). Differences in cDNA inputs were corrected by normalization to *HPRT1* cDNA levels. Relative quantitation of target cDNA was determined by the formula 2^−ΔCT^, with ΔCT denoting fold increase above control. When indicated, an ISG score was calculated as the median of the fold change expression of *RSAD2*, *OAS1*, *Mx1*, *IFI27*, *ISG15*, and *IFI44L*. A list of the TaqMan probes used in this study is supplied in **Supp Table 4**. For mtRNA, levels of mtDNA-encoded transcripts (*MT-CO1, MT-ND6, MT-ND4, MT-ATP6/8, MT-CYTB)* and *SDHA* were quantified by RT-qPCR using Power SYBR Green (Invitrogen) on 12.5 ng/µL cDNA (primers given in **Supp Table 5**). Mt-rRNA *MT-RNR1* and *MT-RNR2*, were quantified by RT-qPCR using Power SYBR Green (Invitrogen) on 0.125 ng/µL cDNA due to their high abundance (primers given in **Supp Table 5**). Analysis was performed as for TaqMan assays.

### Functional validation of CRISPR/Cas9 knockouts

Control, cGAS_KO and STING_KO BJ-5ta cells were stimulated for 24 h with 0.5 µg/ml herring testes synthetic dsDNA > 500 bp (HT-DNA; D6898; Sigma-Aldrich) complexed with Lipofectamine 2000 (Invitrogen), according to the manufacturer’s instructions, or lipofectamine alone. Lack of *IFNB1* mRNA induction upon HT-DNA treatment confirmed cGAS/STING functional impairment in knockout cells compared to controls.

KO of MAVS and MDA5 was validated by transfecting cells with 0.1 µg/mL dsRNA analogue poly(I:C) (tlrl-picwlv, InvivoGen) complexed with lipofectamine (Invitrogen), according to the manufacturer’s instructions, or lipofectamine alone and added to cells for 24 h at 37°C. Lack of *IFNB1* mRNA induction upon poly(I:C) treatment confirmed MDA5 and MAVS functional impairment in KO cells compared to controls.

KO of RIG-I was validated by exposing cells to Sendai virus (SeV, ATCC-VR-907), a potent trigger of RIG-I (Strahle et al., 2006), and confirming lack of *IFNB1* mRNA induction in KO cells relative to controls.

### mtDNA copy number determination

Total DNA from 0.5 × 10^6^ cells was extracted using the Dneasy Blood and Tissue Kit (Qiagen) following the manufacturer’s instructions. DNA concentrations were determined by photometry (Nanodrop) and 15 ng/µL and 7.5 ng/µL DNA were used to perform qPCR of the mtDNA sequences included in *MT-CO2, MT-ND1,* D-loop and the nuclear gene *GAPDH* (primers given in **Supp Table 6**). qPCR was performed using Power SYBR Green (Invitrogen). Ratios of 2^−ΔCT^ for *MT-CO2, MT-ND1,* and D-loop over *GAPDH* of the two DNA concentrations were averaged and expressed as fold of data in control cells.

### mtDNA and mtRNA depletion

mtDNA depletion in BJ-5ta cells was induced by treatment with 100 µM ddC (Sigma-Aldrich) in medium supplemented with uridine 50 µg/mL and sodium pyruvate 1mM, for 10 days before analysis. To control for mtDNA and mtRNA depletion, mtDNA copy number and mtRNA levels were measured as described above.

### Mitochondrial respiration

15 000 cells per well were seeded in quintuplicates in XF96 Cell Culture Microplates 24 h prior to the assay, allowing adherence in complete growth medium at 37°C and 5% CO₂. Prior to measurement, cells were washed and incubated in XF Base Medium supplemented with 10 mM glucose, 2 mM glutamine, and 1 mM pyruvate (pH 7.4) for 1 h at 37°C in a non-CO₂ incubator to equilibrate. The XF96 sensor cartridge was hydrated overnight with XF Calibrant and loaded with stressors: port A (1 µM oligomycin), port B (0.5–2 µM FCCP), port C (0.5 µM rotenone + 0.5 µM antimycin A) for Mito Stress Test; adjusted for ATP Rate Assay as per kit protocol. The assay began with 3 baseline OCR measurements, followed by sequential injections: oligomycin to inhibit ATP synthase, revealing proton leak; FCCP to uncouple and elicit maximal respiration; and rotenone/antimycin A to inhibit electron transport, defining non-mitochondrial OCR. Data were normalized to cell number measured with BCA protein assay or for consistency across wells. Basal respiration was determined as the average OCR before oligomycin injection, subtracting non-mitochondrial OCR (post-rotenone/antimycin A). ATP-linked respiration (mitochondrial ATP production rate) was calculated as basal respiration minus oligomycin-induced OCR, representing the fraction coupled to ATP synthesis. Results were expressed as pmol O₂/min per µg of cells, with statistical analysis on normalized parameters. Wells with OCR <10 pmol/min or high variability (>20% CV across baseline) were excluded. All conditions within an experiment used identical media, seeding, and injection volumes for comparability.

### Immunofluorescence staining and confocal microscopy

BJ-5ta cells were grown on a glass coverslip in a 24-well plate 24 h before fixation. Cells were washed and fixed with 4% paraformaldehyde in PBS and blocked/permeabilized with 0.2% Triton X-100, in 5% bovine serum albumin/PBS for 30 min. Cells were incubated with primary antibody in PBS – 5% BSA buffer (dilution indicated in primary antibody list in **Supp Table 2**) overnight in a humidified chamber at 4°C followed by 3 PBS washes before addition of secondary antibody solution in PBS – 5% BSA (Secondary antibody list in **Supp Table 3**) for 30 min at room temperature. Slides were mounted in Prolong Diamond (ThermoFisher). Single-plane images were acquired with Nikon AX or Leica SP8-SMD confocal microscope using a 63.0×/1.4 oil objective.

### dsRNA specific digestion

dsRNA was specifically digested using ShortCut RNase III (NEB, M0245) in a controlled reaction that cleaves long dsRNA into siRNA-sized fragments (∼18–25 bp) while leaving ssRNA intact (Nicholson & al, 2013). Fixed cells were incubated in 70 µL nuclease-free water, 10 µL 10X ShortCut Reaction Buffer, 10 µL 10X MnCl₂, and 1 µL ShortCut RNase III (1U/µg dsRNA) for 100 µL final volume at 37°C for 20 min. Reactions were stopped by adding 10 µL 100 mM EDTA (pH 8.0) per 100 µL.

### Mitochondrial footprint measurement

Raw images were opened in Fiji and cropped to single cells or defined regions of interest to avoid overlapping structures from neighbouring cells. MiNA plugin applies a standardized preprocessing pipeline consisting of contrast enhancement, background reduction, and smoothing, followed by binarization of the mitochondrial signal to generate a segmented mitochondrial mask. Default MiNA parameters (including thresholding method and ridge detection settings, when applicable) were used unless otherwise specified. Within MiNA output, the “mitochondrial footprint” is defined as the image area (2D) occupied by segmented mitochondrial signal in the binarized mask, expressed in calibrated units (e.g., µm²). To ensure meaningful quantification, only images with well-resolved, clearly labelled mitochondria were included, in line with MiNA recommendations. Samples with weak labelling, high background fluorescence, or poor segmentation (as judged by mismatch between the binary mask and the underlying fluorescence signal) were excluded prior to footprint quantification.

### Mitochondrial membrane potential measurement

Cells were seeded on Ibidi 8 well glass-bottom dishes (IBIDI) suitable for high-resolution live imaging and maintained in medium equilibrated at 37 °C and 5% CO₂. Cells were incubated with TMRE (ThermoFisher) at a final concentration of 100 nM for 30 min at 37 °C to allow dye equilibration across the plasma and mitochondrial membranes. After loading, cells were gently washed once and imaging was performed immediately to minimize dye redistribution and photobleaching. To verify that TMRE fluorescence reflected mitochondrial membrane potential, a subset of samples was treated with the protonophore CCCP (Merck- C2759) 10 µM for 15 min before TMRE loading to depolarize mitochondria and provide a low-potential reference. Live-cell imaging was performed on a Zeiss spinning disk confocal microscope equipped with a 561 nm excitation laser and appropriate emission filter (580–650 nm) for TMRE detection, using an oil immersion objective (63×). Laser power and exposure time were minimized to limit photobleaching and phototoxicity while maintaining sufficient signal-to-noise ratio; these settings were kept constant for all conditions within an experiment. For each field of view, a single optical section was acquired. TMRE images were exported as 16-bit TIFF files and analysed in FIJI/ImageJ. Regions of interest (ROIs) corresponding to mitochondrial mask of individual cells were identified by manual thresholding on the TMRE channel and mean fluorescence intensity per ROI was measured using the “Measure” function. To derive a relative measure of mitochondrial membrane potential, TMRE fluorescence intensity per cell was normalized to the average intensity of untreated controls.

### Sm-FISH

Gene-specific primary probes (up to 24 per transcript, see **Supp Table 7**) were designed using Oligostan software to target optimal sequences with uniform ΔG at 37°C, each containing a common 20 nucleotide FLAP sequence at the 3’ end. Primary probes (100 µM stock in water) were mixed in equimolar proportions and diluted 1/5 in TE buffer. The diluted mixture was pre-hybridized with secondary probes (FLAP-Cy3/Cy5, 100 µM) at 2:1 molar ratio in NEB 3.1 buffer for 3 min at 85°C followed by 3 min at 65°C and 5 min at 25°C on a thermocycler. Cells on #1.5H coverslips were rinsed in PBS, fixed 15 min in 4% PFA/PBS at RT, washed 3× 5 min PBS, then permeabilized overnight at 4°C (or 2 h at RT) in 70% ethanol. Cells were then incubated in 15% formamide (Merck) in SSC buffer (Merck) for 15 min at RT. The cells were then incubated with the duplexed probes conjugated with fluorescent dyes in hybridization buffer (1× SSC, 15% formamide, 20% dextran sulfate, 10µg/L *E. coli* tRNA, 10mg/mL RNase-free BSA, 100 mM VRC) overnight at 37°C in a humid chamber. The next day, cells were washed twice in 15% formamide SSC 30 min at RT. Cells were then washed in PBS three times and mounted in Prolong Diamond. When the protocol was combined with immunostaining, the immunofluorescence protocol was done as described above before the smFISH protocol omitting the permeabilization step in 70% ethanol. Single-plane images were acquired with Nikon AX or Leica SP8-SMD confocal microscope using a 63.0×/1.4 oil objective.

### Quantification of dsRNA signal intensity and dot area

Fixed cells were immunostained for dsRNA (J2 antibody) and TOMM40, and imaged via confocal microscopy as single optical section, as described above. In FIJI, the TOMM40 channel was thresholded to include mitochondrial structures while excluding noise. The thresholded image was converted to a binary mask, then used as ROI and overlaid onto the dsRNA channel. The mean intensity value of dsRNA was measured in mask-covered regions. Mitochondrial dsRNA intensity was reported as mean fluorescence intensity which is the average pixel intensity over the mitochondrial mask surface. The area of the dsRNA particles within the mitochondrial mask was assessed by the *Particle Analysis* tool following manual thresholding and binarization of the dsRNA dots within the mitochondrial mask.

### 3D STED microscopy

Cells cultured on #1.5H glass coverslips were fixed with 4% paraformaldehyde and immunostained with primary antibodies against TOMM40 and dsRNA (J2), followed by secondary antibodies conjugated to Abberior STAR RED (excitation 638 nm/emission 655 nm) for TOMM40 and Abberior STAR Orange (excitation 589 nm/emission 616 nm) for dsRNA, diluted 1:1000 in 5% BSA in PBS buffer for 30 min at Room Temperature (RT). Samples were mounted in Abberior Mount Antifade Solid (medium optimized for STED) and cured 24 h at RT before imaging to ensure refractive index matching (n=1.518). Imaging used the Abberior Facility Line inverted STED microscope with an Olympus 63×/1.42 NA oil-immersion objective (UPLXAPO60XO), deformable mirror module for adaptive optics, and pulsed lasers: 561 nm (STAR Orange), 640 nm (STAR RED), 775 nm STED depletion for both dyes. Time-gated detection (0.5–5 ns window) and pixel dwell time 10–50 µs minimized photobleaching. The signal is accumulated by line 3 times. Z-step was 50 nm for 3D stacks; field of view 1024×1024 pixels (40 nm XY pixel size). 3D deconvolution (built-in Lucy-Richardson, 10–20 iterations) post-processed raw stacks in Imspector software, exporting multichannel TIFFs with calibrated metadata.

### Imaris 3D analysis

3D STED TIFF stacks (TOMM40 and dsRNA) were selected for *Surfaces creation* with absolute intensity threshold optimized to delineate mitochondrial network and identify dsRNA particles. Surface grain size was set to 40 nm, and small TOMM40 positive artifacts not connected to the network (<0.05 µm^3^) as well as too small J2 dots (<0.001 µm^3^) were discarded. DsRNA particles with distance > 100 nm to the mitochondrial network were classified as "outside mitochondria", and the % of cytosolic over the total number of particles was reported. For cytosolic dsRNA spots and mitochondrial ones, *Statistics* tab displayed per-object volume (µm³) directly from voxel count × calibrated voxel dimensions. Sphericity was calculated as ψ = (π^ (1/3) × (6V) ^ (2/3)) / A, where V=volume, A=surface area; values range 0–1 (1=perfect sphere).

### Quantification of cytosolic mtRNA

After BJ-5ta cells of a fully confluent 150 cm^2^ flask (between 6.10^7^ and 12.10^7^ cells) were detached with trypsin, cells were pelleted at 300g for 5 min at 4°C, and washed in PBS twice. The pelleted cells were then resuspended in 500 µL of NaCl 150 mM, HEPES 50 mM and digitonin (50 to 200 µg/mL) in PBS and incubated for 15 min at 4°C under agitation. Cell suspensions were centrifuged again at 1000g for 5 min at 4°C and pellet P1 containing unbroken cells and heavy elements like nuclei was saved. The supernatant was centrifuged at 17 000g for 10 min and pellet P2 containing organelles was saved. The supernatant of P2 was considered as the cytosolic fraction, from which a fraction was saved for western blotting. The rest of the supernatant was mixed 1:1 in volume with RNA lysis buffer from the RNAqueous-Micro Kit (Ambion) and RNA extraction was performed using the standard protocol. mtRNA levels were quantified as described above except that Ct were normalized to *ACTB* and not to *HPRT1*. To compensate for the effect of loss of MTPAP on the abundance of mtRNA transcripts, ratios of 2^−ΔCT^ of cytosolic fractions were normalized to the ones of the total fraction.

### MitoString

Total RNA was isolated from 10^6^ cells using RNeasy Mini Kit with on-column DNase I digestion (Qiagen) to remove DNA contamination. RNA (100 ng) was hybridized with the MitoString CodeSet (55-gene panel adapted from *Wolf and Mootha, 2014*; *Foged et al., 2025*, see **Supp Table 8**) at 67°C for 19 h, then cooled to 45°C before NanoString nCounter processing. Hybridized samples were loaded onto the nCounter Prep Station for automated probe-target purification via magnetic beads, followed by digital counting on the Digital Analyzer. Raw RCC files were processed in nSolver, results were first normalised to the average of housekeeping gene probes. Each transcript abundance in MTPAP_KO or patient fibroblasts was normalised to the average of the respective control cells.

### Transmission electron microscopy

Cells were seeded on 6 wells/plate, fixed with 1.6% glutaraldehyde in 0.1 M cacodylate buffer (pH 7.4) for 2 h, rinsed and postfixed in 1% osmium tetroxide and 1% potassium ferrocyanide in 0.1 M cacodylate buffer before dehydration in ethanol and Epoxy resin embedding. Seventy nanometer sections were contrasted, using a 5% Lanthanide salt solution (a mixture of gadolinium acetate and samarium acetate) for 7 min followed by 6 minutes lead citrate. Observations and pictures were performed with a transmission electron microscope Jeol JEM 1400 equipped with a GATAN RIO9 CMOS camera.

### Quantification and statistical analysis

Statistical analysis was conducted using GraphPad Prism 10 software. Data were displayed as means ± SEM unless otherwise noted. Superplots were tested by paired t-test (2 tail) or one-way ANOVA on the mean of each experiment (Lord et al., 2020). Otherwise, measurements between two groups were performed with Mann-Whitney (one tail) or Wilcoxon (one tail) tests. Groups of three or more were analysed by Kruskal-Wallis as indicated. Grouped analyses were interrogated by two-way ANOVA. Values of n repeats and statistical parameters for each experiment are reported in the figures and figure legends. A p value < 0.05 was considered significant.

## Acknowledgments

The authors thank Claire Pujol, Timothy Wai, and Benedetta Ruzzenente for their valuable comments and advice. The authors are grateful to Alexis Jourdain and Vamsi Mootha for sharing their Mitostring codeset, and Zofia M. Chrzanowska-Lightowlers for providing the fibroblast cell line of P5.

The authors acknowledge Fondation NRJ, Sorbonne Université (Hélios project), Abberior Instruments, ICM.Quant (RRID:SCR_026393) for the STED 3D imaging. Within the Structure Fédérative de Recherche Necker (INSERM US24/CNRS UMS3633), the authors thank the Cell Imaging Core Facility, and Béatrice Durel, for the confocal imaging, as well as the Metabolic Analysis Core Facility and Ivan Nemazanyy, for the Seahorse analysis. The authors acknowledge the Electron Microscopy facility CCMA (Centre Commun de Microscopie Appliquée) from « Université Côte d’Azur », part of the « Microscopie Imagerie Cytométrie d’Azur » GIS IBiSA labeled platform, supported by Université Côte d’Azur, the “Région Sud », the Département 06, CPER, FEDER and Christelle Boscagli for technical help.

Y.J.C. acknowledges the European Research Council (786142-E-T1IFNs), and a state subsidy managed by the National Research Agency (ANR, France) under the ‘Investments for the Future’ programme bearing the reference ANR-10-IAHU-01. Y.J.C. is supported by a UK Medical Research Council Human Genetics Unit core grant (MC_UU_00035/11). A.L. acknowledges funding from ANR (ANR-22-CE15-0012-01) and Inserm (dotation).

## Authors contributions

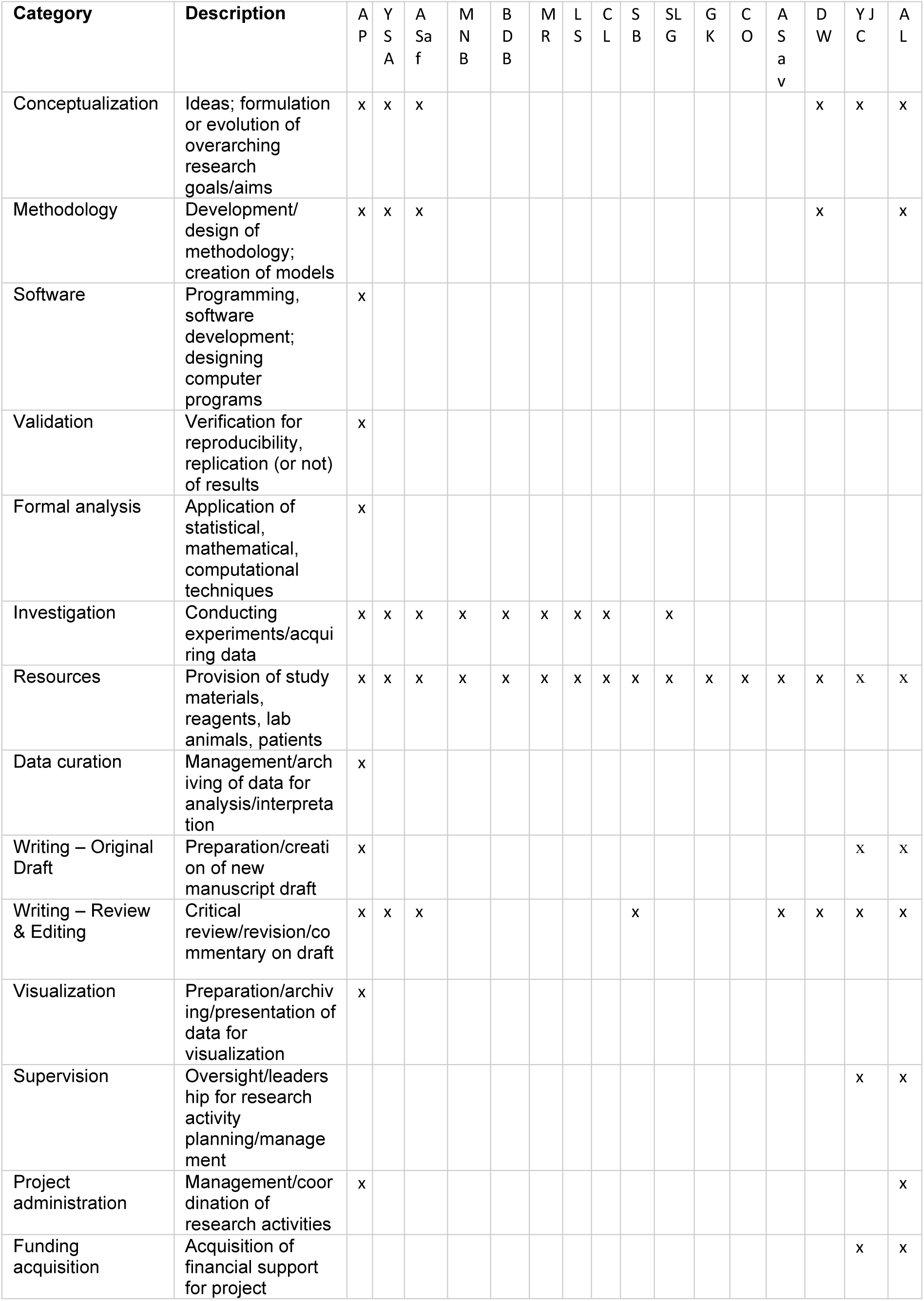

## Supplementary tables

**Table S1.**
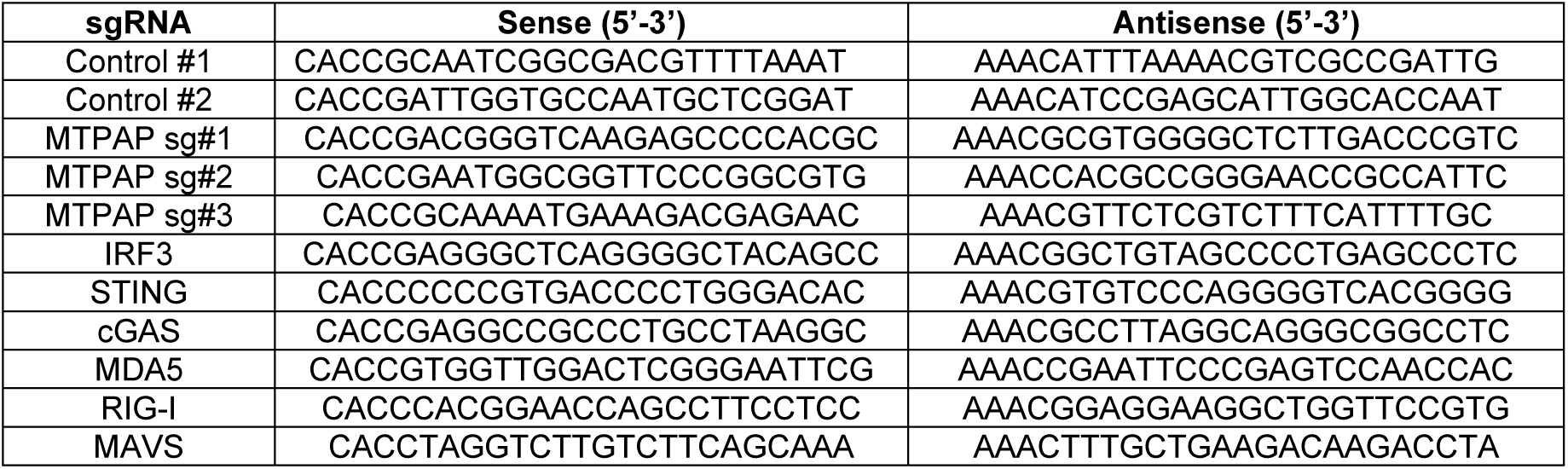
Table of sgRNAs.

**Table S2.** List of primary antibodies used in this study for western blotting and immunofluorescence.

| Antibodies | Host species | Source | Reference | Dilution |
| --- | --- | --- | --- | --- |
| Cofilin | Rabbit monoclonal | Cell Signaling | 5175 | 1:5000 |
| Vinculin | Mouse monoclonal | Santa Cruz | sc-73614 | 1:5000 |
| MTPAP | Mouse monoclonal | Genetex | GTX70156 | 1:1000 |
| Human OXPHOS cocktail | Mouse monoclonal | ThermoFisher | 45-8199 | 1:1000 |
| CYTB | Rabbit polyclonal | Proteintech | 55090-1-AP | 1:1000 |
| IRF3 | Rabbit monoclonal | Cell Signalling | 11904S | 1:1000 |
| STING | Rabbit monoclonal | Cell Signalling | 13647S | 1:1000 |
| cGAS | Rabbit monoclonal | Cell Signalling | 15102S | 1:1000 |
| MAVS | Mouse monoclonal | Santa Cruz | sc-166583 | 1:100 |
| MDA5 | Rabbit monoclonal | Cell Signalling | 5321 | 1:1000 |
| RIG-I | Rabbit monoclonal | Cell Signalling | 3743 | 1:1000 |
| TOMM40 | Rabbit polyclonal | Proteintech | 18409-1-AP | 1:1000 |
| dsRNA (J2) | Mouse monoclonal | Jena Bioscience | RNT-SCI-10010200 | 1 :2000 |
| GRSF1 | Rabbit polyclonal | Merck | HPA036985 | 1 :1000 |
| PNPT1 | Rabbit polyclonal | Abcam | ab96176 | 1 :1000 |
| LAMIN A/C | Mouse monoclonal | Cell Signalling | 4777 | 1 :1000 |
| TIM44 | Rabbit polyclonal | Proteintech | 13589-1-AP | 1:1000 |
| TFAM [C9] | Mouse monoclonal | Santa Cruz | sc-376672 | 1 :100 |

**Table S3.** List of secondary antibodies used in this study for western blotting and immunofluorescence.

| Antibodies | Host species | Source | Reference | Dilution |
| --- | --- | --- | --- | --- |
| IRDye 800CW | Goat anti-rabbit | Li-COR | 926-32211 | 1:10000 |
| IRDye 800CW | Goat anti-mouse | Li-COR | 925-32210 | 1:10000 |
| IRDye 640RD | Goat anti-rabbit | Li-COR | 926-68071 | 1:10000 |
| IRDye 640RD | Goat anti-mouse | Li-COR | 926-68070 | 1:10000 |
| Alexa Fluor 488 | Goat anti-rabbit | ThermoFisher | A-11008 | 1:1000 |
| Alexa Fluor 488 | Goat anti-mouse IgG2a | ThermoFisher | A-21131 | 1:1000 |
| Alexa Fluor 555 | Goat anti-rabbit | ThermoFisher | A-31572 | 1:1000 |
| Abberior STAR ORANGE | Goat anti-mouse | Abberior | STORANGE-1001 | 1 :200 |
| Abberior STAR RED | Goat anti-rabbit | Abberior | STRED-1002 | 1:200 |

**Table S4:** List of Taqman primers used in this study for qPCR.

| Primers | Sequence |
| --- | --- |
| *HPRT1* | Hs03929096_g1 |
| *IFI27* | Hs01086370_m1 |
| *IFI44L* | Hs00199115_m1 |
| *ISG15* | Hs00192713_m1 |
| *RSAD2* | Hs01057264_m1 |
| *IFNB1* | Hs01077958_s1 |
| *OAS1* | Hs00973637_m1 |
| *MX1* | Hs00895608_m1 |

**Table S5.**
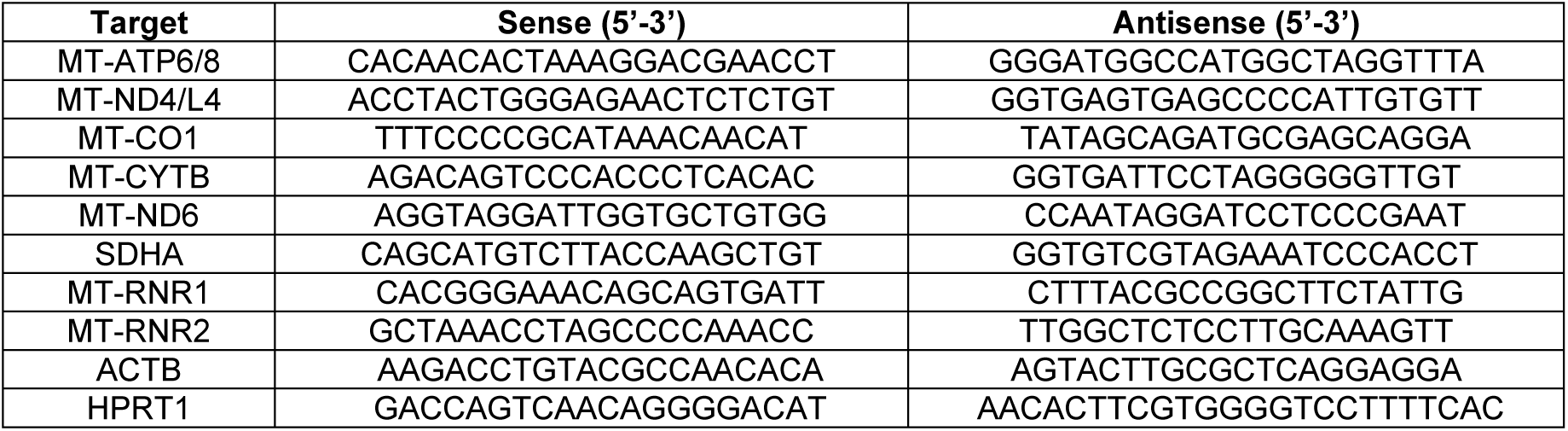
: List of SYBR primers used in this study for qPCR of RNA.

**Table S6.**
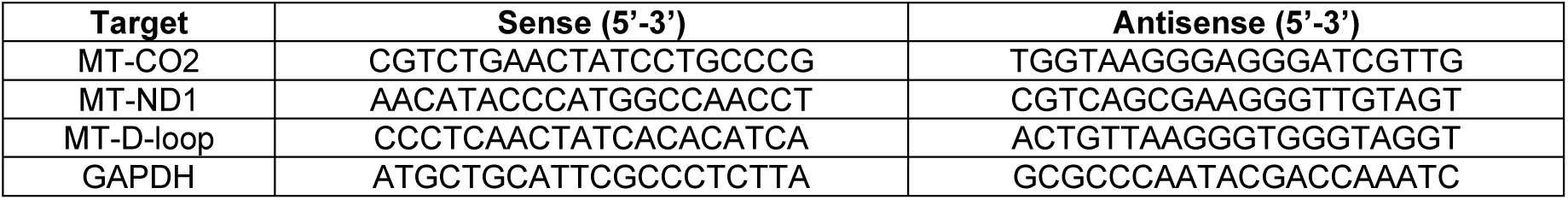
: List of SYBR primers used in this study for qPCR of DNA.

**Table S7.** Sequences of smFISH probes (Excel file)

**Table S8.** Codeset design for Mitostring (Excel file)

**Supplementary data to Figure 1:**
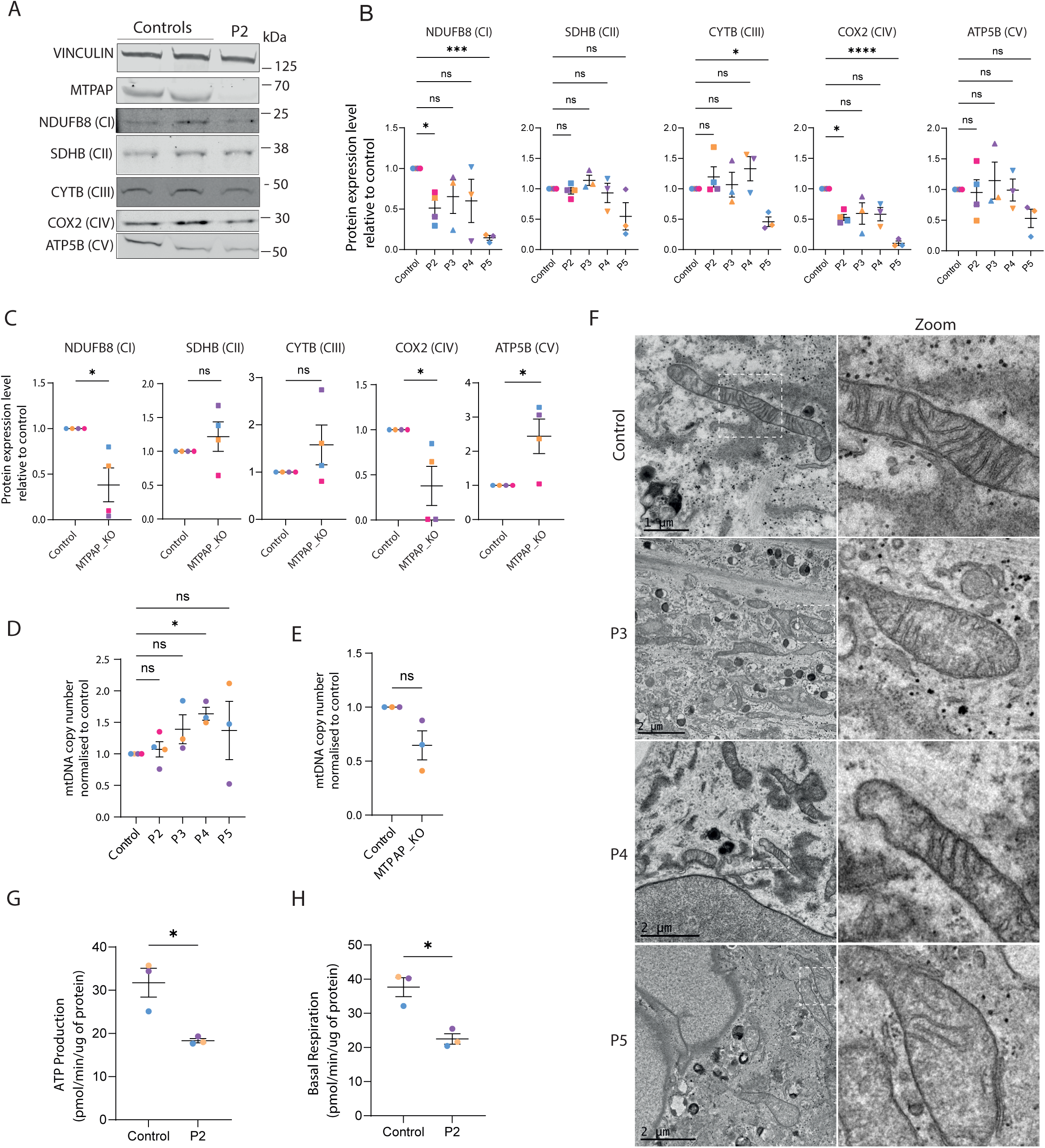
Loss of MTPAP results in defective mitochondrial respiration associated with upregulation of IFN signalling. **A:** Western blot showing levels of MTPAP and mitochondrial protein NDUFB8, SDHB, CYTB, COX2, ATP5B and in lysates of fibroblasts from patient P2 compared to fibroblasts from 2 healthy donor (controls). Vinculin is used as a loading control. Image representative of 3 different experiments. **B-C:** Quantification of mitochondrial protein levels in patient primary fibroblasts (P2, P3, P4 and P5) (**B**) and MTPAP_KO cells (average of sg#1, #2, #3) (**C**) compared to relevant controls. Mean ± SEM; n≥3 experiments, ns indicates non significance, * p< 0.05, *** p<0.001, **** p< 0.0001 in two-way ANOVA with Holm-Sidak multiple comparison test for (**B**) and Mann-Whitney tests for (**C**). **D-E:** Relative levels of mtDNA expressed as the averaged relative levels of *MT-CO2*, *MT-ND1* and mtDNA D-loop signal measured by qPCR in the DNA of patient fibroblasts (**D**) and MTPAP_KO (sg#2) cells (**E**) compared to relevant controls. Mean ± SEM; n≥3 experiments; ns indicates non significance, * p<0.05 in Kruskal-Wallis with Dunn’s comparison test for (**D**) and one sample Wilcoxon tests for (**E**). **F:** Representative TEM images, and zoomed inset, of fibroblasts from patients P3, P4 and P5 and one control showing mitochondrial architecture and cristae. **G-H:** ATP production (**G**) and basal respiration (**H**) measured by Seahorse assay in fibroblasts from P2 versus 3 averaged control cells. Mean ± SEM; n=3 experiments, each colour is a different experiment, * indicates p<0.05 in Mann-Whitney test.

**Supplementary data to Figure 2:**
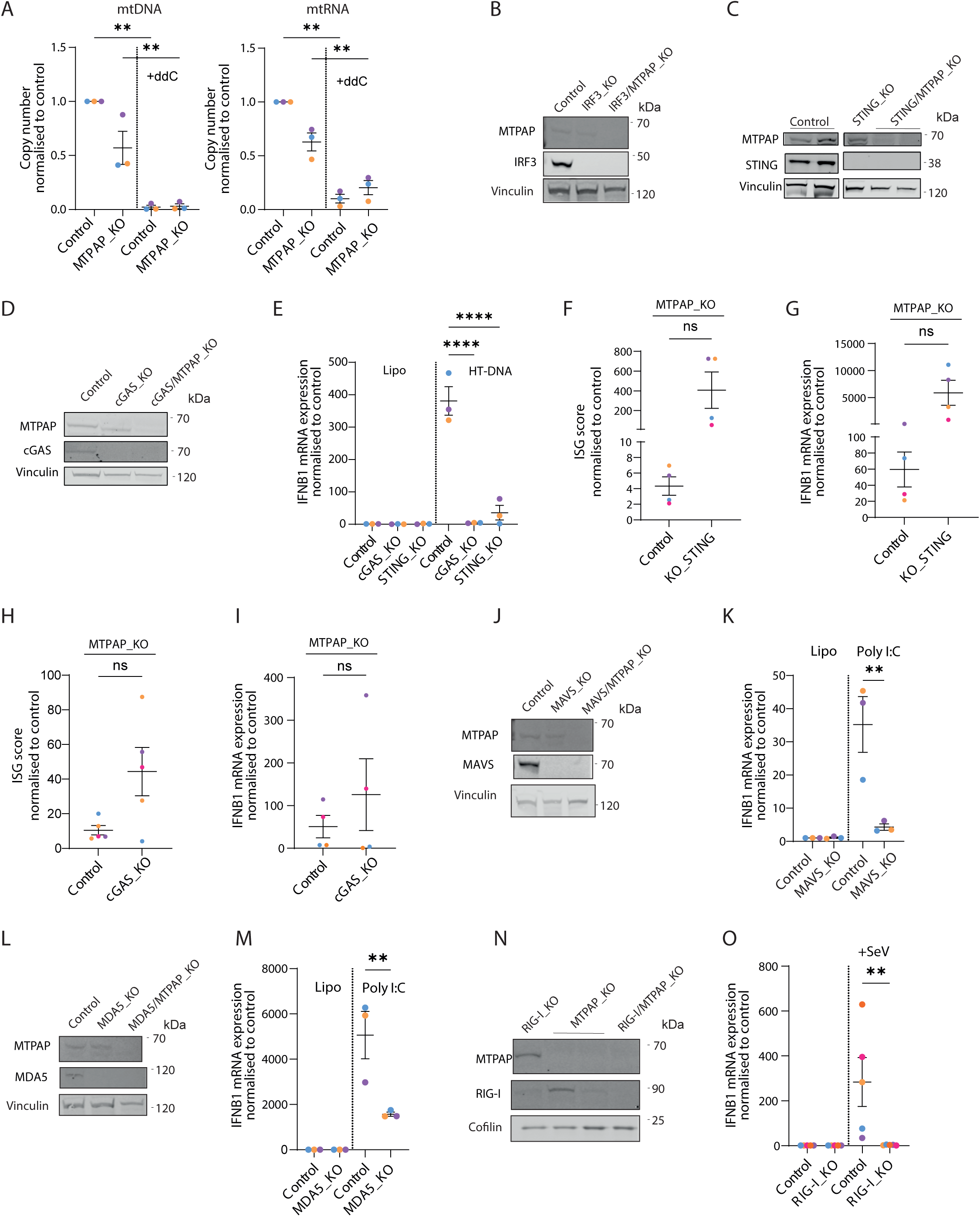
Mitochondrial nucleic acid sensing involves the cytosolic RNA sensor MDA5 and its adaptor MAVS. **A: (Left)** Relative levels of mtDNA expressed as the averaged relative levels of *MT-CO2*, *MT-ND1* and mtDNA D-loop signals and (**Right**) of mtRNA expressed as the averaged relative expression levels of *MT-ATP6/8*, *MT-ND4*, *MT-CO1*, *MT-CYTB* and *MT-ND6* in lysates of control cells and of MTPAP_KO cells (sg#2) treated for 10 days with ddC. Mean ± SEM; n=3 experiments, each colour is a different experiment, ** indicates p<0.01, in two-way ANOVA with Holm-Sidak multiple comparison test. **B-C-D-J-L-N:** Western blot analysis showing protein levels of MTPAP and IRF3 (**B**), STING (**C**), cGAS (**D**), MAVS (**J**), MDA5 (**L**), RIG-I (**N**), in lysates of double KO cells (sg#2 for MTPAP_KO) compared to controls. Vinculin or cofilin are used as loading controls. Pictures representative of 3 different experiments. **E:** mRNA levels of *IFNB1* measured by qPCR in cGAS/MTPAP and STING/MTPAP double KO BJ-5ta cells compared to controls 24 h after stimulation with transfected HT-DNA (sg#2 for MTPAP_KO). Mean ± SEM; n=3 experiments, each colour is a different experiment, **** indicates p< 0.0001 in two-way ANOVA with Holm-Sidak multiple comparison test. **F-H:** ISG score measured by qPCR in BJ-5ta cells double KO for STING/MTPAP (**F**) cGAS/MTPAP (**H**) cells compared to controls (sg#2 for MTPAP_KO). Mean ± SEM; n≥4 experiments, each colour is a different experiment, MTPAP_KO using sg#2; ns indicates non significance for Wilcoxon tests. **G-I:** *IFNB1* mRNA expression measured by qPCR in double KO STING/MTPAP (**G**) cGAS/MTPAP (**I**) cells compared to controls (sg#2 for MTPAP_KO). N=4 experiments, each colour is a different experiment; ns indicates non significance for Wilcoxon tests. **K-M:** *IFNB1*mRNA expression levels in MAVS_KO (**K**) and MDA5_KO (**M**) cells compared to BJ-5ta controls when transfected with poly(I:C). Mean ± SEM; n=3 experiments, ** indicates p<0.01, *** p< 0.001 in two-way ANOVA with Holm-Sidak multiple comparison test. **O:** *IFNB1*mRNA expression levels in RIG-I_KO BJ-5ta cells compared to controls Infected with Sendai Virus. Mean ± SEM; n=5, ** indicates p<0.01 in two-way ANOVA with Holm-Sidak multiple comparison test.

**Supplementary data to Figure 3:**
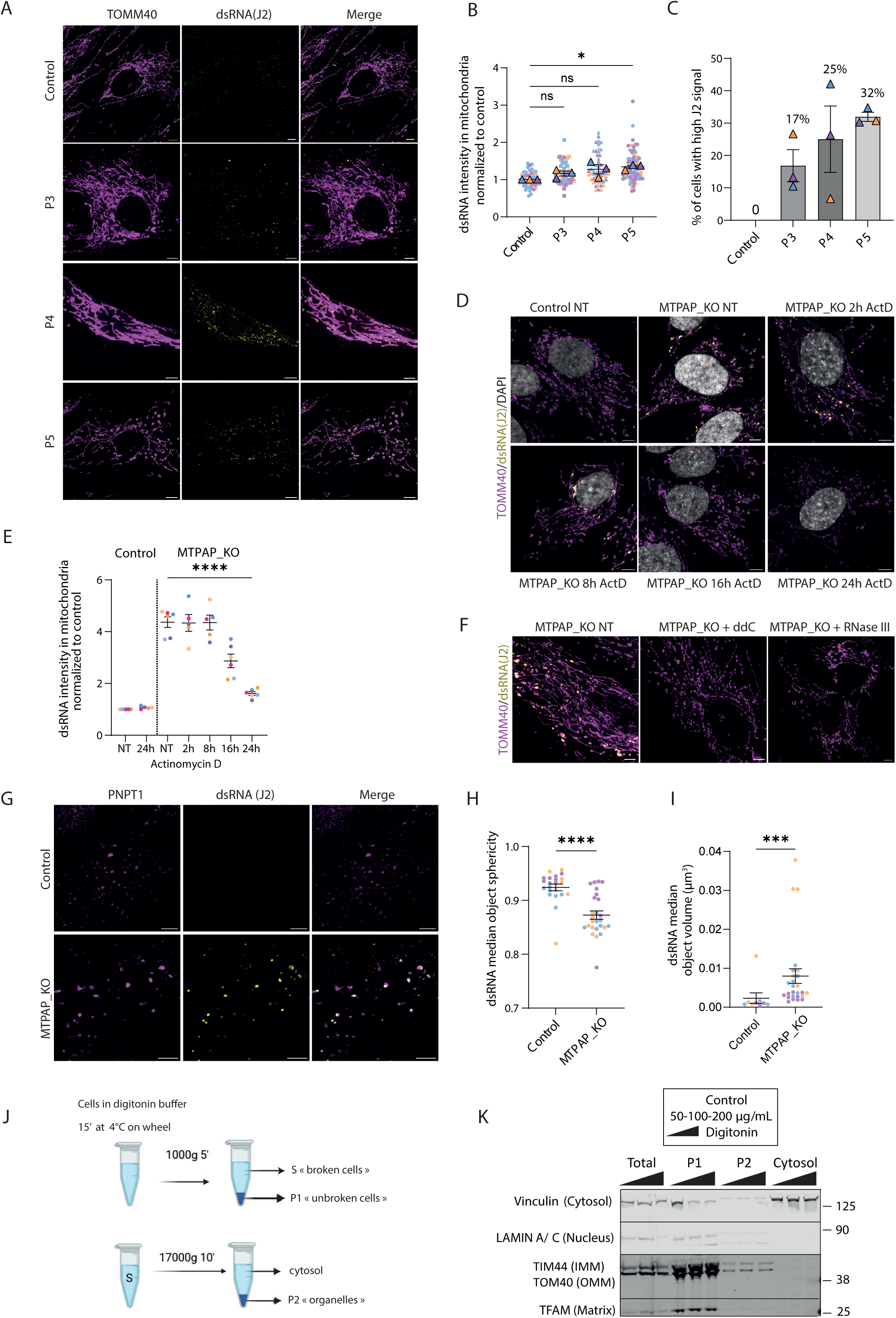
mt-dsRNA are released into the cytosol via mitochondrial pores. **A:** Representative confocal microscopy image of immunostained mitochondrial protein TOMM40 and dsRNA (J2) in control and in patient P3, P4 and P5 fibroblasts. Scale bar: 5µm. **B:** Quantification of average pixel intensity of immunostaining of dsRNA signal in mitochondria in control cells and in patients’ cells (P3, P4 and P5). Superplot, mean ± SEM; n=3 experiments; each borderless point represents the measure of a cell; each colour is a different experiment; ns indicates non significance, * p<0.05 in one-way ANOVA with Dunnett multiple comparison test test performed on the mean value for each experiment **C:** Percentage of cells with a J2 signal 1.5-fold above the average of 3 controls. Mean ± SEM; n=3 experiments, each colour is a different experiment. **D:** Representative confocal microscopy image of immunostained TOMM40 and dsRNA (J2) in control and MTPAP_KO BJ-5ta cells (sg#2) untreated (NT) or treated with actinomycin D (ActD) for up to 24h. Scale bar: 5 µm. **E:** Quantification of pixel intensity of immunostaining of dsRNA signal in mitochondria in control and MTPAP_KO cells (average of sg#1, #2, #3) untreated (NT) and treated with actinomycin D overtime. Mean ± SEM; n=6 experiments, **** indicates p<0.0001 in two-way ANOVA with Holm-Sidak multiple comparison test. **F:** Representative confocal microscopy image of immunostained TOMM40 and dsRNA (J2) in MTPAP_KO cells (sg#2) non treated (NT), treated in culture for 10 days with ddC, or untreated in culture but treated after fixation with dsRNA specific RNase III (see Methods). Scale bar: 5 µm. **G:** Representative confocal microscopy image of immunostained PNPT1 and dsRNA (J2) in control and MTPAP_KO cells (sg#3). Scale bar: 5µm. **H-I:** Quantification of the median sphericity (**H**) and median volume (**I**) of dsRNA particles in control and MTPAP_KO cells (sg#2). Mean ± SEM; n=3, each dot represents a different cell, each colour represents a different experiment, *** indicates p<0.001, **** p<0.0001 in Mann-Whitney test. **J:** Schematic representation of the experimental set-up to isolate the cytosolic fraction using digitonin and differential centrifugation. **K:** Western blot analysis of the different fractions generated to isolate cytosolic fractions. Vinculin is used as a cytosolic marker, Lamin A/C as a nuclear fraction marker, TIM44 and TOM40 as mitochondrial membrane markers and TFAM as mitochondrial matrix marker. Note the absence of all markers but vinculin in the cytosolic fraction.

**Supplementary data to Figure 4:**
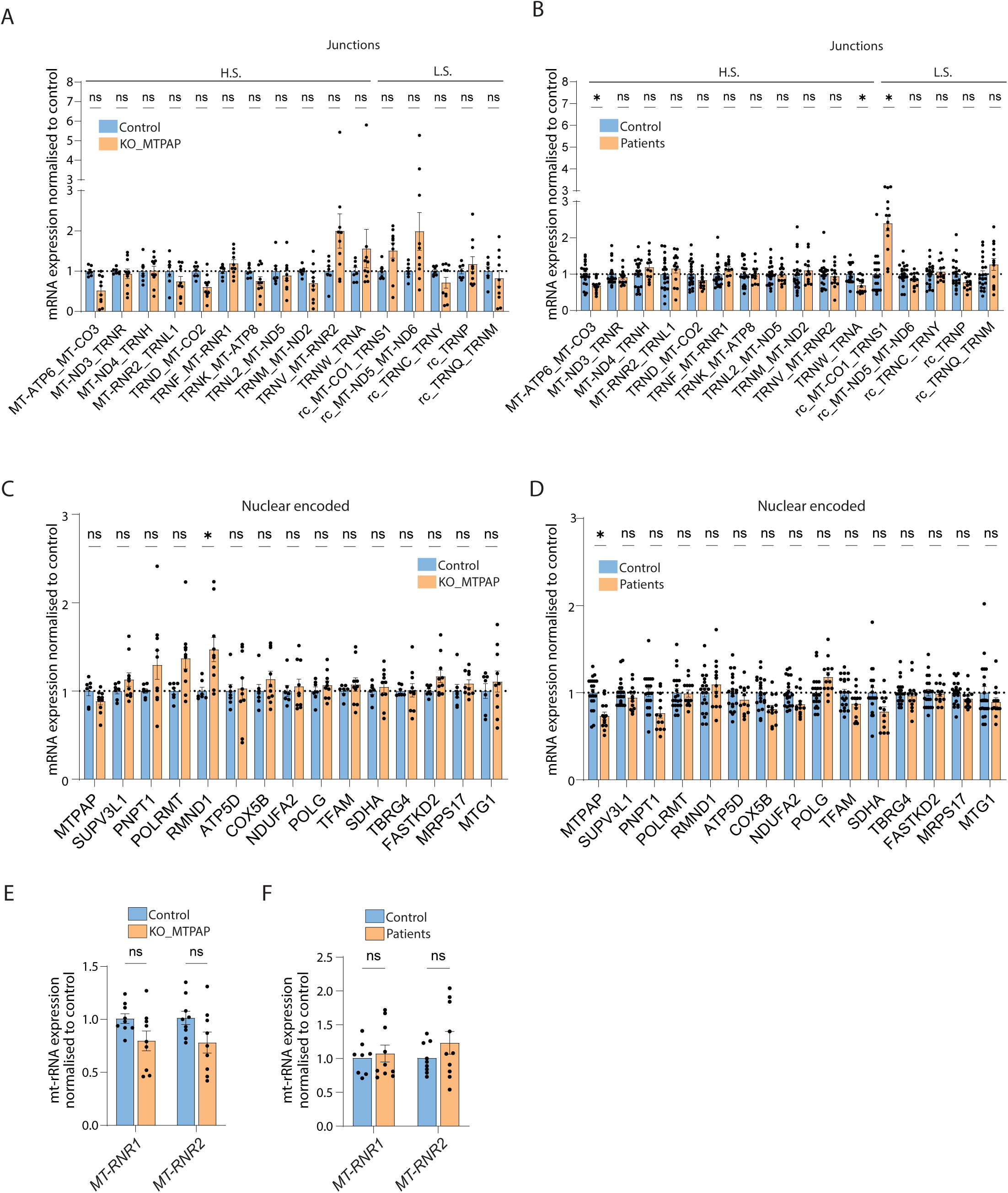
MTPAP loss causes loss of mt-mRNA in parallel with accumulation of non-coding mtRNA. **A-B:** Relative levels of indicated junction mtRNAs in lysates of MTPAP_KO cells (sg#1, #2, #3 each collected 3 times) (**A**) and fibroblasts from patients with *MTPAP* mutations (P2, P3, P4, P5 each collected 3 times) (**B**) compared to respective controls and quantified using MitoString. H.S.: heavy strand, L.S.: light strand. Mean ± SEM; n=3 experiments; each point corresponds to a different sample, ns indicates non significance, * p<0.05 in multiple Mann-Whitney with Holm-Sidak comparison tests. **C-D:** Relative levels of nuclear encoded mRNA coding for proteins involved in mtRNA metabolism in lysates of MTPAP_KO cells (sg#1, #2, #3 each collected 3 times) (**A**) and fibroblasts from patients with *MTPAP* mutations (P2, P3, P4, P5 each collected 3 times) (**B**) compared to respective controls and quantified using MitoString. Mean ± SEM; n=3 experiments; each point corresponds to a different sample, ns indicates non significance, * p<0.05 in multiple Mann-Whitney with Holm-Sidak comparison tests. **E-F:** Expression levels of mt-rRNA *MT-RNR1* and *MT-RNR2* measured by qPCR in MTPAP_KO (sg#1, #2, #3 each collected 3 times) and in MTPAP patients (P3, P4, P5 each collected 3 times) (**F**) cells compared to control. Mean ± SEM; n=3 experiments; each point corresponds to a different sample; ns indicates non significance in Mann-Whitney test.

## References

Abrisch, R. G., Gumbin, S. C., Wisniewski, B. T., Lackner, L. L. & Voeltz, G. K. Fission and fusion machineries converge at ER contact sites to regulate mitochondrial morphology. J Cell Biol 219, e201911122 (2020).

Al-Shamsi, A., Hertecant, J. L., Souid, A.-K. & Al-Jasmi, F. A. Whole exome sequencing diagnosis of inborn errors of metabolism and other disorders in United Arab Emirates. Orphanet J Rare Dis 11, 94 (2016).

Antonicka, H., Sasarman, F., Nishimura, T., Paupe, V. & Shoubridge, E. A. The Mitochondrial RNA-Binding Protein GRSF1 Localizes to RNA Granules and Is Required for Posttranscriptional Mitochondrial Gene Expression. Cell Metabolism 17, 386–398 (2013).

Begeman, A., Smolka, J. A., Shami, A., Waingankar, T. P. & Lewis, S. C. Spatial analysis of mitochondrial gene expression reveals dynamic translation hubs and remodeling in stress. Science Advances 11, eads6830 (2025).

Borowski, L. S., Dziembowski, A., Hejnowicz, M. S., Stepien, P. P. & Szczesny, R. J. Human mitochondrial RNA decay mediated by PNPase–hSuv3 complex takes place in distinct foci. Nucleic Acids Res 41, 1223–1240 (2013).

Brown, T. A. & Clayton, D. A. Release of replication termination controls mitochondrial DNA copy number after depletion with 2′,3′-dideoxycytidine. Nucleic Acids Res 30, 2004–2010 (2002).

Bryant, J. D., Lei, Y., VanPortfliet, J. J., Winters, A. D. & West, A. P. Assessing Mitochondrial DNA Release into the Cytosol and Subsequent Activation of Innate Immune-related Pathways in Mammalian Cells. Current Protocols 2, e372 (2022).

Chen, W. W., Freinkman, E., Wang, T., Birsoy, K. & Sabatini, D. M. Absolute Quantification of Matrix Metabolites Reveals the Dynamics of Mitochondrial Metabolism. Cell 166, 1324–1337.e11 (2016).

Chrzanowska-Lightowlers, Z. M. & Lightowlers, R. N. Mitochondrial RNA maturation. RNA Biology 21, 1065–1076 (2024).

Chujo, T. et al. LRPPRC/SLIRP suppresses PNPase-mediated mRNA decay and promotes polyadenylation in human mitochondria. Nucleic Acids Res 40, 8033–8047 (2012).

Crosby, A. H. et al. Defective Mitochondrial mRNA Maturation Is Associated with Spastic Ataxia. The American Journal of Human Genetics 87, 655–660 (2010).

Crow, Y. J. & Casanova, J.-L. Human life within a narrow range: The lethal ups and downs of type I interferons. Science Immunology 9, eadm8185 (2024).

Dauletbaev, N., Cammisano, M., Herscovitch, K. & Lands, L. C. Stimulation of the RIG-I/MAVS Pathway by Polyinosinic:Polycytidylic Acid Upregulates IFN-β in Airway Epithelial Cells with Minimal Costimulation of IL-8. J Immunol 195, 2829–2841 (2015).

David, C. et al. Gain-of-function human UNC93B1 variants cause systemic lupus erythematosus and chilblain lupus. J Exp Med 221, e20232066 (2024).

Dhir, A. et al. Mitochondrial double-stranded RNA triggers antiviral signalling in humans. Nature 560, 238–242 (2018).

Dunker, W. et al. TDP-43 prevents endogenous RNAs from triggering a lethal RIG-I-dependent interferon response. Cell Reports 35, 108976 (2021).

Epstein, J., Veera, S., Pappas, J. & Shah, B. C. 7238 A rare cause of primary ovarian failure in an adolescent due to a genetic mutation in mitochondrial poly-A-polymerase (mTPAP) mRNA. J Endocr Soc 8, bvae163.1398 (2024).

Eyck, L. V. et al. Biallelic Mutations in MTPAP Associated with a Lethal Encephalopathy. Neuropediatrics 51, 178–184 (2020).

Fiedler, M., Rossmanith, W., Wahle, E. & Rammelt, C. Mitochondrial poly(A) polymerase is involved in tRNA repair. Nucleic Acids Res 43, 9937–9949 (2015).

Foged, M. M. et al. Cytosolic N6AMT1-dependent translation supports mitochondrial RNA processing. Proceedings of the National Academy of Sciences 121, e2414187121 (2024).

Genin, E. C. et al. CHCHD10 mutations promote loss of mitochondrial cristae junctions with impaired mitochondrial genome maintenance and inhibition of apoptosis. EMBO Mol Med 8, 58–72 (2016).

Gerencser, A. A., Mookerjee, S. A., Jastroch, M. & Brand, M. D. Measurement of the Absolute Magnitude and Time Courses of Mitochondrial Membrane Potential in Primary and Clonal Pancreatic Beta-Cells. PLOS ONE 11, e0159199 (2016).

Gustafsson, C. M., Falkenberg, M. & Larsson, N.-G. Maintenance and Expression of Mammalian Mitochondrial DNA. Annu. Rev. Biochem. 85, 133–160 (2016).

Hansen, K. G., Baxter-Koenigs, A., Weiss, C. A., McShane, E. & Churchman, L. S. Transcription arrest induces formation of RNA granules in mitochondria. Life Sci. Alliance 8, e202403082 (2025).

Hensen, F. et al. Mitochondrial RNA granules are critically dependent on mtDNA replication factors Twinkle and mtSSB. Nucleic Acids Res 47, 3680–3698 (2019).

Hiramatsu, Y. et al. Complex hereditary peripheral neuropathies caused by novel variants in mitochondrial-related nuclear genes. J Neurol 269, 4129–4140 (2022).

Hooftman, A. et al. Macrophage fumarate hydratase restrains mtRNA-mediated interferon production. Nature 615, 490–498 (2023).

Hou, F. et al. MAVS Forms Functional Prion-like Aggregates to Activate and Propagate Antiviral Innate Immune Response. Cell 146, 448–461 (2011).

Iborra, F. J., Kimura, H. & Cook, P. R. The functional organization of mitochondrial genomes in human cells. BMC Biol 2, 9 (2004).

Idiiatullina, E. et al. Heterozygous de novo dominant negative mutation of REXO2 results in interferonopathy. Nat Commun 15, 6685 (2024).

Jourdain, A. A. et al. GRSF1 Regulates RNA Processing in Mitochondrial RNA Granules. Cell Metabolism 17, 399–410 (2013).

Killarney, S. T. et al. Executioner caspases restrict mitochondrial RNA-driven Type I IFN induction during chemotherapy-induced apoptosis. Nat Commun 14, 1399 (2023).

Kim, J. et al. VDAC oligomers form mitochondrial pores to release mtDNA fragments and promote lupus-like disease. Science 366, 1531–1536 (2019).

Kim, S. et al. Mitochondrial double-stranded RNAs govern the stress response in chondrocytes to promote osteoarthritis development. Cell Reports 40, 111178 (2022).

Kim, S. et al. RNA 5-methylcytosine marks mitochondrial double-stranded RNAs for degradation and cytosolic release. Molecular Cell 84, 2935–2948.e7 (2024).

Krieger, M. R. et al. Trafficking of mitochondrial double-stranded RNA from mitochondria to the cytosol. Life Science Alliance 7, (2024).

Lepelley, A. et al. Enhanced cGAS-STING–dependent interferon signaling associated with mutations in ATAD3A. J Exp Med 218, e20201560 (2021).

Lepelley, A., Wai, T. & Crow, Y. J. Mitochondrial Nucleic Acid as a Driver of Pathogenic Type I Interferon Induction in Mendelian Disease. Front. Immunol. 12, (2021).

Lord, S. J., Velle, K. B., Mullins, R. D. & Fritz-Laylin, L. K. SuperPlots: Communicating reproducibility and variability in cell biology. J Cell Biol 219, e202001064 (2020).

Mai, N., Chrzanowska-Lightowlers, Z. M. A. & Lightowlers, R. N. The process of mammalian mitochondrial protein synthesis. Cell Tissue Res 367, 5–20 (2017).

McShane, E. et al. A kinetic dichotomy between mitochondrial and nuclear gene expression processes. Molecular Cell 84, 1541–1555.e11 (2024).

Nagaike, T., Suzuki, T., Katoh, T. & Ueda, T. Human Mitochondrial mRNAs Are Stabilized with Polyadenylation Regulated by Mitochondria-specific Poly(A) Polymerase and Polynucleotide Phosphorylase*. Journal of Biological Chemistry 280, 19721–19727 (2005).

Nicholson, A. W. Ribonuclease III mechanisms of double-stranded RNA cleavage. WIREs RNA 5, 31–48 (2014).

Pajak, A. et al. Defects of mitochondrial RNA turnover lead to the accumulation of double-stranded RNA in vivo. PLOS Genetics 15, e1008240 (2019).

Pearce, S. F. et al. Maturation of selected human mitochondrial tRNAs requires deadenylation. eLife 6, e27596 (2017).

Rai, P. & Fessler, M. B. Mechanisms and effects of activation of innate immunity by mitochondrial nucleic acids. Int Immunol 37, 133–142 (2025).

Ravanbod, M. et al. Clinical and molecular assessment of a spastic ataxia 4 (SPAX4) patient with a novel variant in the MTPAP gene, and a systematic review. Gene 956, 149463 (2025).

Rehwinkel, J. & Gack, M. U. RIG-I-like receptors: their regulation and roles in RNA sensing. Nat Rev Immunol 20, 537–551 (2020).

Rey, T. et al. Mitochondrial RNA granules are fluid condensates positioned by membrane dynamics. Nat Cell Biol 22, 1180–1186 (2020).

Rice, G. I. et al. Assessment of interferon-related biomarkers in Aicardi-Goutières syndrome associated with mutations in TREX1, RNASEH2A, RNASEH2B, RNASEH2C, SAMHD1, and ADAR: a case-control study. The Lancet Neurology 12, 1159–1169 (2013).

Ruzzenente, B. et al. LRPPRC is necessary for polyadenylation and coordination of translation of mitochondrial mRNAs. EMBO J 31, 443–456 (2012).

Ryan, C. M. & Read, L. K. UTP-dependent turnover of Trypanosoma brucei mitochondrial mRNA requires UTP polymerization and involves the RET1 TUTase. RNA 11, 763–773 (2005).

Safieddine, A. et al. HT-smFISH: a cost-effective and flexible workflow for high-throughput single-molecule RNA imaging. Nat Protoc 18, 157–187 (2023).

Sanjana, N. E., Shalem, O. & Zhang, F. Improved vectors and genome-wide libraries for CRISPR screening. Nat Methods 11, 783–784 (2014).

Schonborn, J. et al. Monoclonal antibodies to double-stranded RNA as probes of RNA structure in crude nucleic acid extracts. Nucleic Acids Res 19, 2993–3000 (1991).

Souza, P. V. S. de, et al. New genetic causes for complex hereditary spastic paraplegia. Journal of the Neurological Sciences 379, 283–292 (2017).

Stewart, S. A. et al. Lentivirus-delivered stable gene silencing by RNAi in primary cells. RNA 9, 493–501 (2003).

Strahle, L., Garcin, D. & Kolakofsky, D. Sendai virus defective-interfering genomes and the activation of interferon-beta. Virology 351, 101–111 (2006).

Stringer, B. W. et al. A reference collection of patient-derived cell line and xenograft models of proneural, classical and mesenchymal glioblastoma. Sci Rep 9, 4902 (2019).

Sun, L., Wu, J., Du, F., Chen, X. & Chen, Z. J. Cyclic GMP-AMP Synthase Is a Cytosolic DNA Sensor That Activates the Type I Interferon Pathway. Science 339, 786–791 (2013).

Suomalainen, A. & Nunnari, J. Mitochondria at the crossroads of health and disease. Cell 187, 2601–2627 (2024).

Szczesny, R. J. et al. Human mitochondrial RNA turnover caught in flagranti: involvement of hSuv3p helicase in RNA surveillance. Nucleic Acids Res 38, 279–298 (2010).

Tomecki, R., Dmochowska, A., Gewartowski, K., Dziembowski, A. & Stepien, P. P. Identification of a novel human nuclear-encoded mitochondrial poly(A) polymerase. Nucleic Acids Res 32, 6001–6014 (2004).

Toompuu, M. et al. Polyadenylation and degradation of structurally abnormal mitochondrial tRNAs in human cells. Nucleic Acids Res 46, 5209–5226 (2018).

Tsanov, N. et al. smiFISH and FISH-quant – a flexible single RNA detection approach with super-resolution capability. Nucleic Acids Res 44, e165 (2016).

Valente, A. J., Maddalena, L. A., Robb, E. L., Moradi, F. & Stuart, J. A. A simple ImageJ macro tool for analyzing mitochondrial network morphology in mammalian cell culture. Acta Histochemica 119, 315–326 (2017).

van Esveld, S. L. et al. Mitochondrial RNA processing defect caused by a SUPV3L1 mutation in two siblings with a novel neurodegenerative syndrome. Journal of Inherited Metabolic Disease 45, 292–307 (2022).

Wang, Y. et al. Escaping from CRISPR–Cas-mediated knockout: the facts, mechanisms, and applications. Cell Mol Biol Lett 29, 48 (2024).

Weber, F., Wagner, V., Rasmussen, S. B., Hartmann, R. & Paludan, S. R. Double-Stranded RNA Is Produced by Positive-Strand RNA Viruses and DNA Viruses but Not in Detectable Amounts by Negative-Strand RNA Viruses. Journal of Virology 80, 5059–5064 (2006).

Wilson, W. C. et al. A human mitochondrial poly(A) polymerase mutation reveals the complexities of post-transcriptional mitochondrial gene expression. Hum Mol Genet 23, 6345–6355 (2014).

Wolf, A. R. & Mootha, V. K. Functional Genomic Analysis of Human Mitochondrial RNA Processing. Cell Reports 7, 918–931 (2014).

Xavier, V. J. & Martinou, J.-C. RNA Granules in the Mitochondria and Their Organization under Mitochondrial Stresses. International Journal of Molecular Sciences 22, 9502 (2021).

Xavier, V. et al. Mitochondrial double-stranded RNA homeostasis depends on cell-cycle progression. Life Science Alliance 7, (2024).

Xiao, T. S. & Fitzgerald, K. A. The cGAS-STING pathway for DNA sensing. Mol Cell 51, 135–139 (2013).

Yoon, J. et al. Diminished SUV3 expression and its functional implications in the IFN-enriched monocyte subset of childhood Sjögren’s disease. Rheumatology (Oxford) 64, 4393–4403 (2025).

Ysebrant de Lendonck, L., Martinet, V. & Goriely, S. Interferon regulatory factor 3 in adaptive immune responses. Cell. Mol. Life Sci. 71, 3873–3883 (2014).

